# Metabolic plasticity enables cancer adaptation to acidic tumor ecosystems

**DOI:** 10.64898/2026.01.14.699600

**Authors:** Raafat Chalar, Naheel Khatri, Jowana Obeid, Joon-Hyun Song, Yujie Xiao, Saiful Samad, Andrew Chen, Andrew Resnick, Khadijeh Karbalaei, Janet J. Allopenna, Cungui Mao, Christopher Clarke, Fabiola Velazquez, Bo Chen, Daniel Canal, Yusuf Hannun, Mehdi Damaghi

## Abstract

Cell state plasticity enables cancer cells to rapidly adapt to fluctuating microenvironments without requiring genetic alteration, shaping tumor evolution under stress. Extracellular acidosis is a persistent feature of solid tumors that impose strong selective pressure, yet how cancer cells maintain fitness under acute and chronic acidic conditions remains unclear. Here, we show that adaptation to acidosis is mediated by plastic rewiring of sphingolipid metabolism centered on ceramide turnover. Spatial multi-omics analysis of three-dimensional tumor models revealed enrichment of ceramides within acidic niches, consistent with a stress-induced phenotype. While acute acidosis promoted ceramide accumulation and reduced fitness, chronic exposure selected for cells capable of dynamically redistributing sphingolipid flux across multiple clearance pathways. Functional perturbation demonstrated that inhibition of individual pathways was insufficient to compromise survival, whereas simultaneous disruption of all ceramide clearance routes resulted in cell death, revealing a degenerate metabolic architecture. This network-level flexibility enables cancer cells to maintain fitness by switching between alternative metabolic states under acidic stress. Together, our findings identify sphingolipid metabolic plasticity as an adaptive strategy that supports tumor persistence in acidic ecosystems and suggest that targeting metabolic flexibility, rather than individual pathways, may provide a more effective therapeutic approach.

## Introduction

Cell state plasticity enables cancer cells to rapidly adapt to fluctuating microenvironmental conditions without requiring genetic alteration, providing a key mechanism for survival under stress (*1–4*). Unlike mutation-driven evolution, plasticity allows reversible transitions between phenotypic states, thereby influencing selection dynamics and long-term tumor evolution (*2*, *5–8*). Among these adaptive processes, metabolic plasticity plays a central role by enabling cells to reprogram energy production and biosynthetic pathways in response to microenvironmental constraints such as hypoxia, nutrient limitation, and acidosis (*5*, *9*). Extracellular acidosis is a defining and persistent feature of solid tumors, arising early during tumorigenesis and maintained throughout progression. In ductal carcinoma in situ (DCIS), diffusion-limited oxygen availability promotes glycolytic metabolism and lactic acid accumulation, generating spatially structured acidic niches (*10*). Importantly, acidosis is not restricted to hypoxic regions but is reinforced by aerobic glycolysis, establishing a chronic selective pressure within tumor ecosystems. Survival in these environments therefore requires adaptive strategies that operate across both spatial and temporal scales (*5*, *11*).

Lipids, and in particular sphingolipids, are central regulators of cellular fate, integrating structural, metabolic, and signaling functions (*12–14*)(*15*). Ceramides promote stress responses and apoptosis, whereas downstream metabolites such as sphingosine-1-phosphate and glycosphingolipids support proliferation and survival (*15*) (*16*). While this balance has been described as a rheostat controlling cell fate, it remains unclear how sphingolipid metabolism is dynamically regulated within spatially heterogeneous tumor microenvironments, and whether it contributes to adaptive plasticity under chronic stress (*13*, *17*).

Despite growing interest in tumor metabolomics, most studies lack spatial resolution and fail to capture how metabolic processes are organized within ecological tumor habitats (*7*, *18–20*). Moreover, how metabolic networks adapt under persistent acidosis, and whether such adaptation reflects pathway-specific dependencies or broader network-level properties, remains poorly understood.

Here, we integrate spatial multi-omics, functional perturbation, and evolutionary analysis to investigate sphingolipid metabolism in acidic tumor ecosystems. We identify ceramide turnover as a central adaptive process and demonstrate that cancer cells maintain fitness under acidosis not by committing to a single metabolic pathway, but by exploiting a plastic and degenerate network of ceramide clearance routes. This flexibility enables dynamic rerouting of metabolic flux and supports survival under chronic stress. Furthermore, single-nuclei multi-omic analysis suggests that these state transitions are coordinated by distributed transcriptional programs likely mediated by chromatin-dependent regulation rather than a single dominant regulator.

## Results

### Ecosystem profiling using spatial multi-omics of 3D spheroids identifies sphingolipids in acidic habitats

Ductal carcinoma of the breast experiences localized acute and chronic acidosis in the tumor ecosystem during early carcinogenesis (Extended fig. 1a)(*9*). To resolve the spatial organization of acidic tumor habitats, we applied an integrated multi-omics approach combining MALDI imaging mass spectrometry (MALDI-MSI) and multiplex immunofluorescence (mIF) in 3D MCF7 spheroids (Fig. 1a, Extended Fig. 1c-e, Extended Fig. 2). These models recapitulate physiologically relevant gradients of oxygen and pH, enabling spatial mapping of metabolic and phenotypic states. Co-registration of MALDI-derived metabolite and lipid profiles with mIF-defined cellular phenotypes revealed distinct metabolic habitats across the spheroid. MALDI-MSI across nine spheroid sections detected 366 ions at 50% FDR (METASPACE) (*23*), of which 52 high-confidence ions (10% FDR) were retained for analysis (Supplementary Table S1).

**Figure 1:**
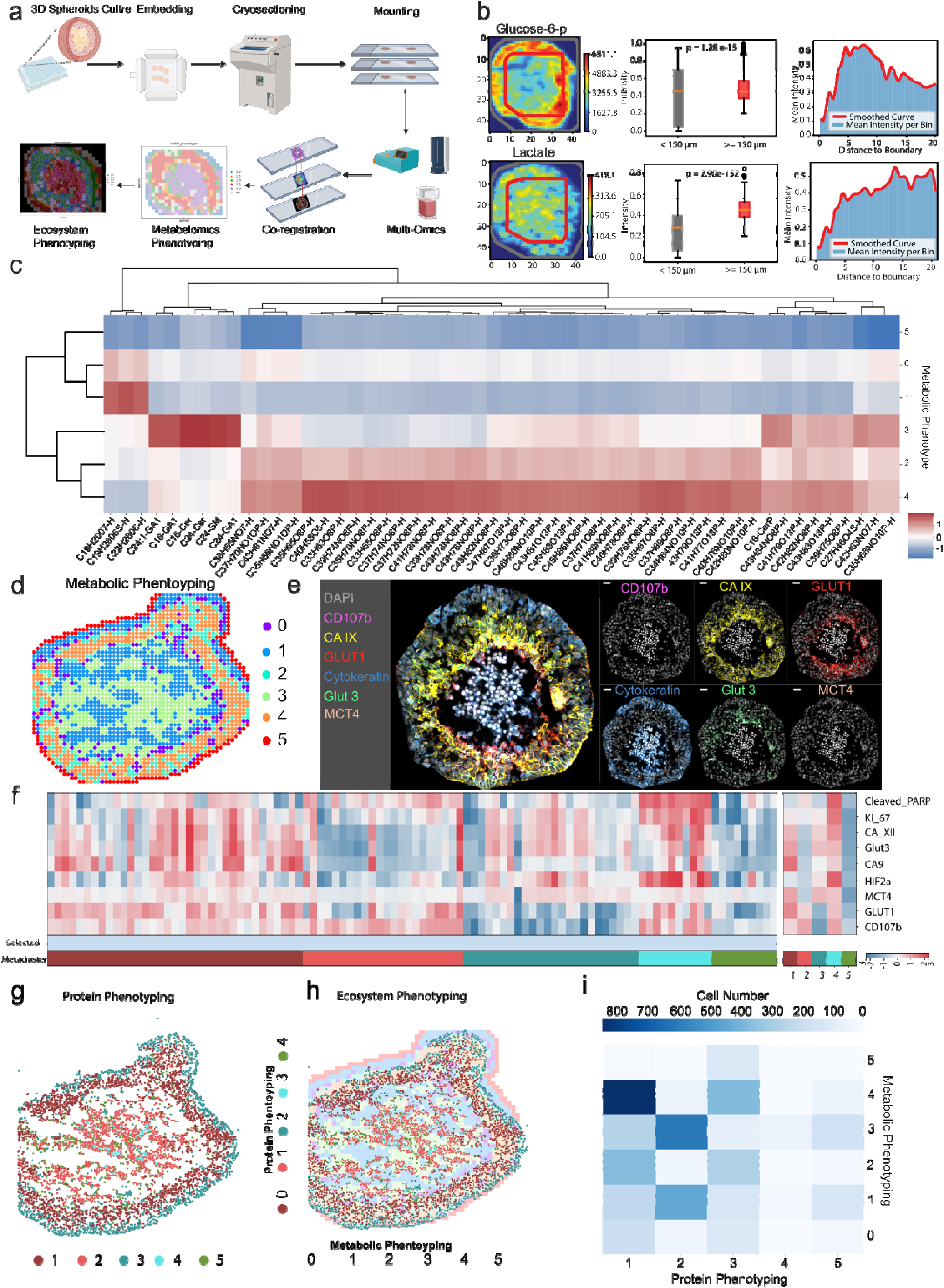
Spatial multi-omics analysis reveals enrichment of ceramides in acidic niche. **a**, Schematic of the experimental workflow, including 3D spheroid culture, embedding, cryo-sectioning, and spatial multi-omics integration, including MALDI imaging, H&E staining, and multiplex mIF. **b**, Spatial intensity maps of glucose-6-phosphate (G6P) and lactate. Boxplots quantify levels in normoxic (0-150 µm from the edge) vs. hypoxic regions (>150 µm from the edge). Statistical analysis was performed using an unpaired t-test. Histogram measures molecule average intensity as a function of the distance from the spheroid boundary. **c**, Heatmap representing the intensity of metabolites and lipids across distinct clusters normalized to scale. **d**, Unsupervised metabolic clustering visualizing spatially resolved metabolic phenotypes. **e**, Multiplex-immunofluorescence (mIF) imaging using MACSima visualizing microenvironmental markers (e.g., CD107b (LAMP2b), CA IX, Glut1, Glut3, cytokeratin) to define phenotypic niches. **f**, Single-cell resolution heatmap of mIF clusters. Scale represents normalized signal intensities. **g**, Unsupervised clustering to identify the spatial organization of the proteomic phenotypes. **h**, Ecosystem phenotyping by co-registering metabolic phenotypes and the protein phenotypes spatial clustering. **i**, Co-occurrence matrix depicting the spatial relationship between metabolic and proteomic phenotypes. Rows correspond to metabolic phenotypes; columns represent proteomic phenotypes. The color scale indicates the number of single cells from protein phenotyping clusters that are localized within the corresponding metabolic phenotyping cluster pixels.

Habitat stratification based on oxygen diffusion distance confirmed expected metabolic organization, with glucose-6-phosphate enriched in normoxic regions (<150 μm from the edge) and lactate in hypoxic cores (Fig. 1b), validating the spatial framework. Unsupervised clustering of MALDI signals segmented the spheroids into distinct metabolic subtypes with layered spatial organization (Fig. 1c and 1d; Extended Fig. 3a). Notably, sphingolipids, including C16- and C24-ceramides, sphingomyelin, and GA1, were enriched in cluster 3, corresponding to the central acidic region, whereas phospholipids (PS, PE, PC) were enriched in outer, proliferative regions (cluster 4) (Extended Fig. 3d).

In parallel, mIF profiling resolved single-cell phenotypic heterogeneity across spheroids (Fig. 1e–g; Extended Fig. 3b and 3c). Integration of MALDI and mIF datasets through spatial co-registration enabled mapping of metabolic states to cellular phenotypes (Fig. 1h and 1i; Extended Fig. 2a). Cells occupying acidic habitats (MACSima phenotype 2; Fig. 1f) co-localized predominantly with MALDI cluster 3, linking acidosis-associated phenotypes to sphingolipid-enriched regions. Targeted sphingolipid MALDI analysis further confirmed elevated ceramides and downstream metabolites within these regions, particularly in spheroid cores (Extended Fig. 2b and 3e). Together, these data establish a spatially resolved association between sphingolipid metabolism and acidic tumor habitats.

### Ceramides are increased in response to acute acidic stress

To determine how acidosis alters sphingolipid metabolism, we exposed MCF7 breast cancer cells to acute acidic conditions (48 hours, pH 6.5; non-adapted, NA) and quantified sphingolipids by LC/MS (Fig. 2a). Acute acidosis led to a significant increase in multiple ceramide species compared to physiological pH (Fig. 2b), indicating that acidic stress induces rapid accumulation of pro-apoptotic sphingolipids. To assess whether this response is spatially recapitulated in tumor-like architectures, we analyzed ceramide distribution in MCF7 spheroids using MALDI imaging. Consistent with the 2D measurements, ceramides were enriched in the spheroid core, corresponding to hypoxic and acidic regions, relative to the normoxic periphery (Fig. 2c). This spatial pattern closely mirrors the metabolic gradients observed in our ecosystem profiling (Fig. 1), confirming that ceramide accumulation is a localized feature of acidic tumor habitats. This response was conserved across additional breast cancer cell lines, including T47D and MDA-MB-231, whereas non-transformed MCF10A cells exhibited an opposite trend (Extended Fig. 4), suggesting a cancer-specific adaptation to acidic stress. Together, these results demonstrate that acute acidosis induces a robust and spatially organized accumulation of ceramides, establishing a stress-associated metabolic state that precedes adaptive remodeling of sphingolipid flux.

**Figure 2.**
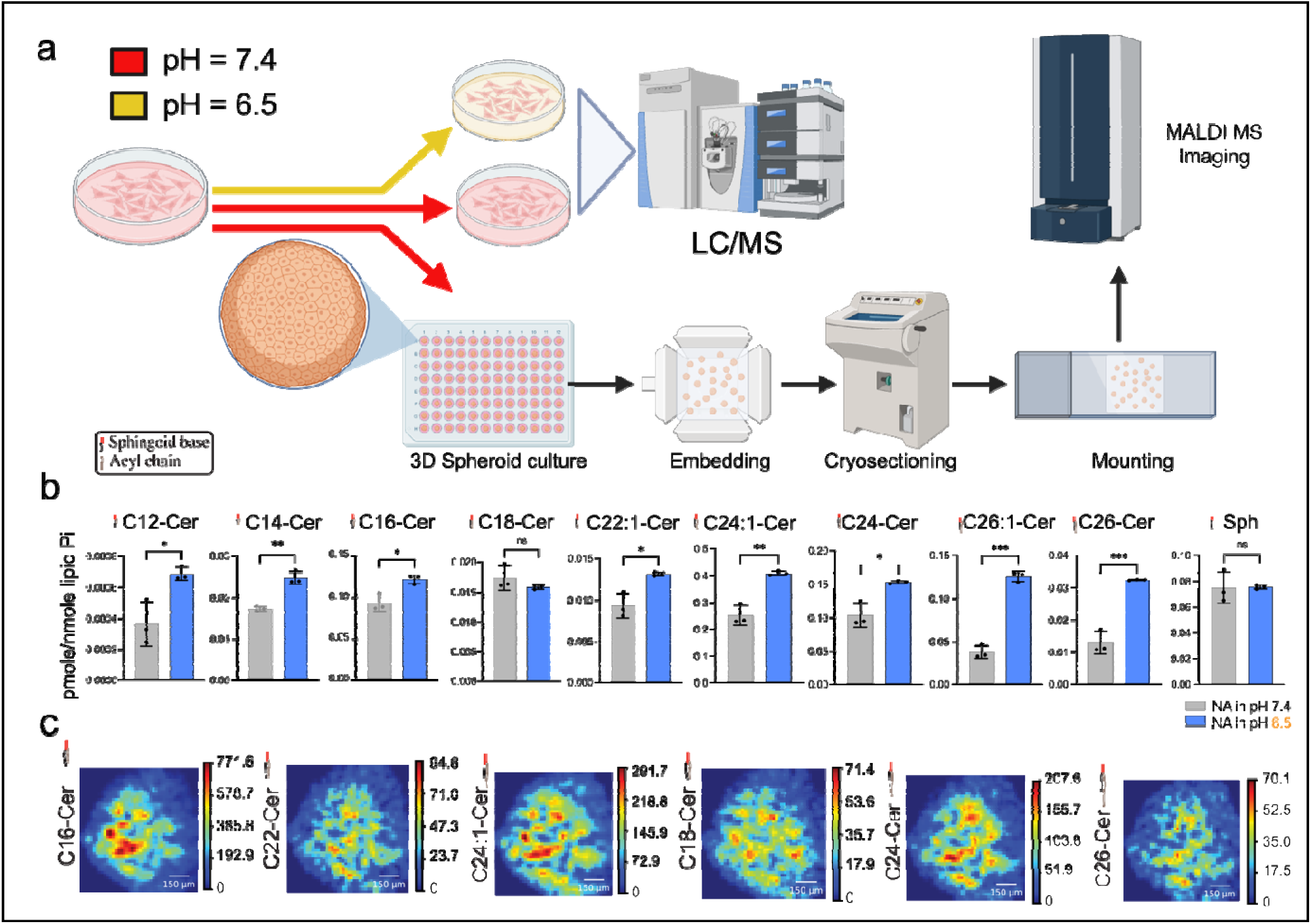
Acute acidosis induces ceramide accumulation in breast cancer cells. **a**, Experimental workflow for sphingolipid profiling in MCF7 cells exposed to physiological (pH 7.4) or acidic (pH 6.5) conditions. Cells were analyzed by LC/MS for bulk lipid quantification and by MALDI mass spectrometry imaging in 3D spheroids following embedding and cryosectioning. **b**, LC/MS quantification of ceramide species in MCF7 cells under physiological and acidic conditions, showing significant increases in multiple ceramide species following acute acidosis (48 h). Data are presented as pmole/nmole lipid Pi. Statistical significance was determined by unpaired t-test (*P < 0.05, **P < 0.01, ***P < 0.001). **c**, MALDI imaging of ceramide species in MCF7 spheroids, demonstrating spatial enrichment of ceramides in the spheroid core corresponding to hypoxic and acidic regions. Scale bars, 150 μm.

### CRISPR-Cas screening reveals cancer cells exploit all SL clearance path to increase fitness in acidic microenvironments

To investigate these sphingolipid metabolic phenotypes in the acidic TME, we employed a CRISPR/Cas synthetic lethality screen on MCF7 cancer cells under physiological pH and acidic pH using a library targeting only genes involved in SL metabolism (Fig. 3a, Supplementary table S2). We identifie key genes critical for cell survival under acidosis (Fig. 3b), including Alkaline Ceramidase-3 (ACER3), Alkaline Ceramidase-2 (ACER2) and Acid Ceramidase (ASAH) - enzymes responsible for conv rting ceramide to sphingosine - as well as Sphingosine Kinase 1 (SK1), which catalyzes the phosphorylation of Sphingosine (Sph) to produce Sphingosine-1-phosphate (S1P). Also, COL4A3BP (encoding the CERT protein) was identified as a top hit. It was shown in other cancers that Ceramide/S1P Axis is an adaptive mechanism in acute acidic microenvironments (*25*). Therefore, we first looked at this pathway, notably Sphingosine Kinase 1 (SK1), Alkaline Ceramidase-1 (ACER1), Alkaline Ceramidase-2 (ACER2) and Alkaline Ceramidase-3 (ACER3) – all enzymes responsible for clearing Cer by converting it to sphingosine (Sph) which can be subsequently metabolized to Sphingosine-1-phosphate (S1P). Cer/S1P axis can also play a pivotal role in determining cellular fate by converting tumor-suppressing ceramides and sphingosine into S1P, a tumor-promoting molecule that activates oncogenic pathways through binding to S1PR (Fig. 3c) (*26*)(*27*). Thus, we performed RT-PCR for the ceramidase enzymes (ACER1, ACER2, ACER3, and ASAH1) as well as the S1P receptor (S1PR3) and found significant increased expression of ACER1, ACER2, ASAH1, and S1PR3 under acute acidosis (Fig. 3d). To assess the effect of the S1P path on spheroid growth, we inhibited SK1 using a specific S1P synthesis inhibitor PF-543 (PF). Surprisingly, inhibiting SK1 did not affect the growth rate of spheroids (Fig. 3e). These unexpected results suggested that cancer cells may have switched to other pathways of ceramide clearance when the S1P path is blocked and possess some degree of metabolic plasticity, enabling them to bypass SK1 dependency. To test this hypothesis, we analyzed our RNA sequencing data, focusing on genes involved in sphingolipid metabolism (Fig. 3f). Differential expression analysis using DESeq2 (*28*), comparing MCF7 cells cultured under acidic versus physiological pH revealed a significant upregulation of UGT8, a key enzyme in the glycosphingolipid (GSL) synthesis pathway. UGT8 was also identified in the CRISPR screen, although with minimal effect compared to ACERs. To assess the role of this pathway, we inhibited GSL synthesis using Eliglustat (El) alone and in combination with PF. Notably, inhibition of GSL synthesis, either alone or together with S1P pathway blockade, did not produce a significant reduction in spheroid growth (Fig. 3g).

**Figure 3.**
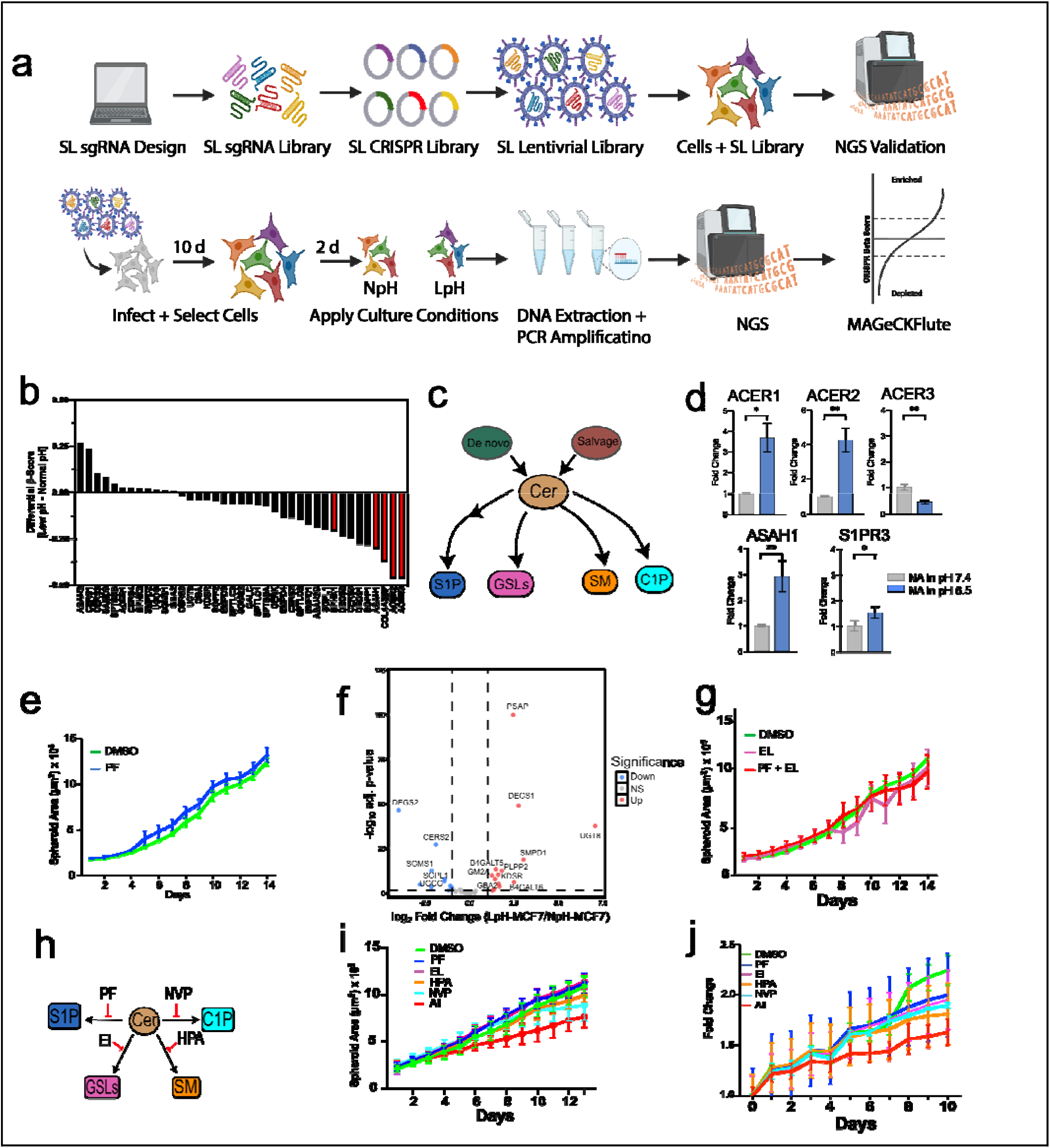
CRISPR screening reveals several genes from pathways of SL metabolism enhance cancer cell fitness under acute acidosis. **a**, Schematic representation of CRISPR/Cas9 screening targeting sphingolipid (SL) metabolism genes. **b**, Differential β-score analysis highlighting genes essential for survival under physiological pH (7.4) and acidic pH (6.5) conditions. **c**, Schematic of sphingolipid metabolism highlighting two ceramide synthesis pathways and four clearance routes. **d**, qRT-PCR analysis of genes involved in the Cer/S1P pathway, including *ACER1-3*, *ASAH1*, *S1PR2-3*, and *SGPL1.* Data are presented as fold changes. Statistical analysis was performed using an unpaired t-test (*P < 0.05, **P < 0.01, ***P < 0.001, ****P < 0.0001). **e**, Growth kinetics of MCF7 spheroids treated with SK1 inhibitor (PF) and vehicle demonstrating no difference. **f**, Volcano plot of sphingolipid metabolism–related genes differentially expressed between MCF7 cells in acid and physiological pH, highlighting coordinated transcriptional rewiring of ceramide-processing pathways. The genes set for the sphingolipid metabolism pathway was retrieved from KEGG hsa00600(*31*, *32*). Results are summarized as log2 fold changes, MCF7 under NpH and LpH, with adjusted p-values using the Benjamini-Hochberg procedure. **g**, Growth curves of MCF7 cells under pharmacologic perturbation of two sphingolipid metabolism using PF and Eliglustat, indicating no differential sensitivity in treated spheroids even in combination of the two drugs. **h**, Model summarizing metabolic all pathways of flux away from ceramides toward pro-survival sphingolipid species. **i**, Proliferation of MCF7 spheroids treated individually and all together with inhibitors targeting distinct sphingolipid metabolic pathways, revealing pathway-specific dependencies. **j**, Longitudinal growth analysis of patient-derived tumor organoids treated with similar inhibitors in i, demonstrating conserved sensitivity to disruption of ceramide clearance in patient derived organoids.

So far, we targeted two major Cer clearance pathways out of four (Fig. 3h), yet cancer cell fitness remained unaffected. This resilience led us to question the extent of metabolic plasticity in cancer cells and how dynamically they can reprogram SL metabolism to maintain their fitness despite disruptions in key clearance pathways. However, to first understand whether cancer cells survive if we block all possible Cers metabolic exit pathways, we simultaneously targeted all ceramide exit pathways together in both spheroids and breast cancer patients derived organoids (PDO) (Fig. 3i and j). To perform this, we included HPA-12, which targets Cer trafficking and mainly disrupts Sphingomyelin (SM) biosynthesis (*29*), and NVP-231, which targets Ceramide conversion to Ceramide-1-phosphate (C1P) (*30*). We found that simultaneous inhibition of all pathways leads to cancer cell death, whereas inhibition of any single one of the four pathways is insufficient to compromise cell viability (Fig. 3i and j). These observations raised key questions: To what extent does each ceramide clearance pathway contribute to cancer cell fitness? And do all pathways provide equivalent fitness advantages, or do some confer a greater benefit? To address these questions, we forced cancer cells to depend on a single Cer metabolic route and assessed their fitness (Extended Fig. 5a). We measured spheroid growth using inhibitors in combinations designed to restrict cells to a single route (Extended Fig. 5b). Our results revealed that selection for the S1P pathway under E+H+N combination conferred the lowest fitness, whereas the SM pathway, particularly under the P+E+N combination, restored spheroid growth, confirming our previous observations (Extended Fig. 5c).

### Cancer cells exhibit remarkable plasticity in regulating sphingolipid metabolism

To assess whether sphingolipid metabolic plasticity enables compensatory pathway switching, we inhibited individual ceramide clearance pathways in spheroids/PDO and performed MALDI imaging to map the resulting reorganization of sphingolipid metabolism (Fig. 4a, Extended Fig. 6, Supplementary video 1). We first evaluated the potency of the inhibitors in suppressing their respective SL products (e.g., HPA targeting SM synthesis, Eliglustat targeting GSL synthesis). Significant reductions in the target metabolic nodes were observed for each inhibitor (Fig. 4a). PF and NVP potency were not evaluated due to the low cellular levels of their targets, S1P and C1P, which were below the detection threshold in MALDI (Extended Figure 2b). We then applied our habitat analysis to various SL species including Cers, SMs, and GSLs (Extended Fig. 8). Notably, treatment with PF or Eliglustat led to increased C16-ceramide accumulation in the inner regions of spheroids, indicating that the Cer/S1P and Cer/GSL pathways are likely major ceramide clearance routes in acidic niches. In contrast, treatment with HPA or NVP reduced ceramide levels in both the inner and outer regions of spheroids, indicating that inhibition of sphingomyelin or C1P synthesis does not lead to ceramide accumulation in these regions. This may reflect compensatory redistribution of sphingolipid flux through alternative metabolic routes, consistent with sphingolipid metabolic plasticity. (Fig. 4a, Extended Fig. 6a). However, inhibition of the Cer/SM pathway with HPA induced a compensatory shift toward the Cer/GSL pathway, as indicated by the pronounced accumulation of HexCer in the outer regions of the spheroids. (Fig. 4b, Extended Fig. 6b). Additionally, more complex GSLs, such as C16-LacCer and C24-Gb3 were also elevated in the inner regions of the spheroids following HPA treatment (Extended Fig. 8). SMs were also impacted by the reprogramming of SL metabolism. HPA treatment effectively reduced SM levels throughout the spheroids (Fig. 4a, Extended Fig. 6c). Notably, suppression of C1P synthesis with NVP-231 treatment had a significant impact on C16-SM levels, particularly in the outer layers of the spheroids.

**Figure 4.**
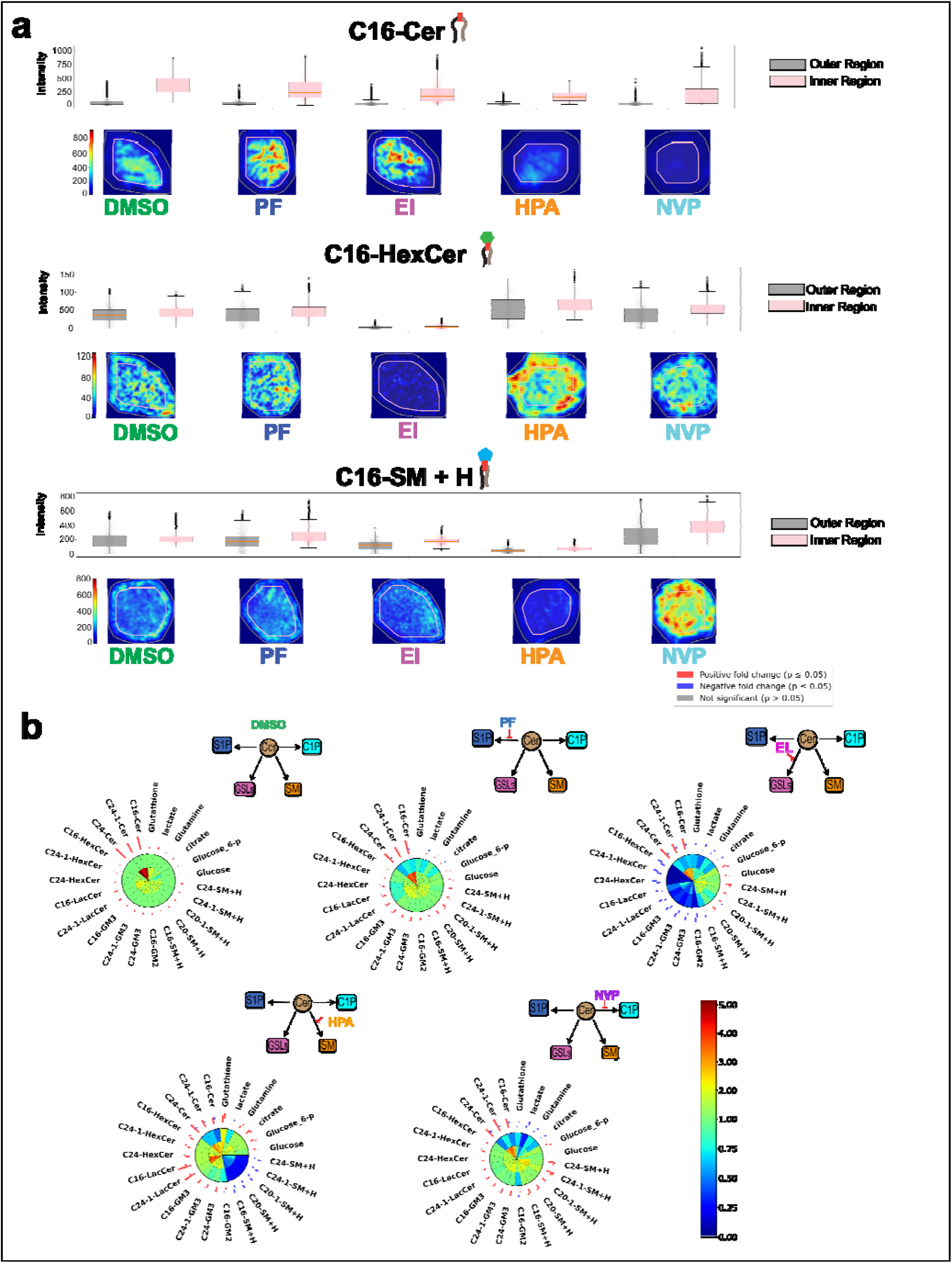
Spatial analysis of phenotypic plasticity and sphingolipids reprogramming in breast cancer spheroids models. **a-c**, MALDI mass spectrometry imaging (MALDI-MSI) of MCF7 spheroids treated with SL inhibitors, highlighting spatial distribution and intensity of selected molecules. Box plots indicate average molecular intensities in inner and outer layers, defined by a 125 µm threshold from the spheroid boundary. **d**, Spatial Phenotype Heatmap for Ecosystem Representation (SPHERE) plots illustrating the spatial response of various molecules to SL inhibitors. Positive and negative fold changes are indicated (P ≤ 0.05). Schematic overlays illustrating the inferred redistribution of ceramide flux into distinct sphingolipid pathways, including S1P, glycosphingolipids (GSLs), sphingomyelin (SM), and ceramide-1-phosphate (C1P), under selective pathway inhibition in PDOs. Collectively, these analyses demonstrate heterogeneous yet coordinated sphingolipid plasticity in patient-derived tumor organoids, supporting ceramide clearance as a conserved adaptive strategy in both complex spheroid and organoid 3D tumor ecosystems.

Given the profound plasticity and compensatory rewiring observed following these pharmacologic perturbations, we developed SPHERE (Spatial Phenotype Heatmap for Ecosystem Representation) plots to visualize the spatial effects of each treatment on several SL species at once. These plots compare the intensity of each molecule post-treatment relative to the outer layer of DMSO controls across two spheroid regions: the outer layers (oxygen-rich, nonacidic) and the inner layers (hypoxic, acidic) (**Figure 4b, Extended Figure 6d**). The SPHERE plots revealed the remarkable plasticity of sphingolipid metabolism, as shown by the distinct spatial metabolic patterns induced by each inhibitor. Interestingly, perturbation of individual ceramide turnover pathways did not yield isolated metabolite changes but instead triggered broader network-level redistribution across sphingolipid classes. HPA most clearly induced compensatory rerouting toward the glycosphingolipid branch, whereas PF, EL, and NVP produced distinct patterns of sphingolipid remodeling across spatial regions of the spheroid. These findings support a model in which ceramide metabolism is highly interconnected, such that inhibition of one pathway generates ripple effects throughout the broader sphingolipid network.

### Ceramide clearance is maintained as a stable adaptation to chronic acidosis

Our acute perturbation studies revealed that although ceramide clearance is broadly required for survival, singular metabolic routes used are less important, pointing to substantial sphingolipid metabolic plasticity. This prompted us to ask whether such flexibility reflects a transient response to acute stress or a stable adaptation to chronic acidosis. Because acidity is a persistent feature of the TME, this distinction is biologically important. Ceramide accumulation is cytotoxic and reduces cellular fitness; therefore, if ceramide clearance is adaptive rather than merely protective, prolonged acid exposure should favor sustained depletion of intracellular ceramides together with their redirection into downstream pro-survival metabolites, including S1P and GSLs. To test this, we adapted MCF7 cells from the same parental population used in Figures 1 and 2 to pH 6.5 for more than three months (Fig. 5a), followed by mass spectrometry analysis as described for Figure 2. Acid-adapted (AA) cells exhibited ceramide levels that were lower than non-adapted (NA) MCF7 cells under acidic condition and even lower or comparable to those observed in physiological pH (Fig. 5b), supporting the conclusion that sustained ceramide clearance constitutes an adaptive strategy under chronic acidic stress rewiring Cer metabolism.

**Figure 5.**
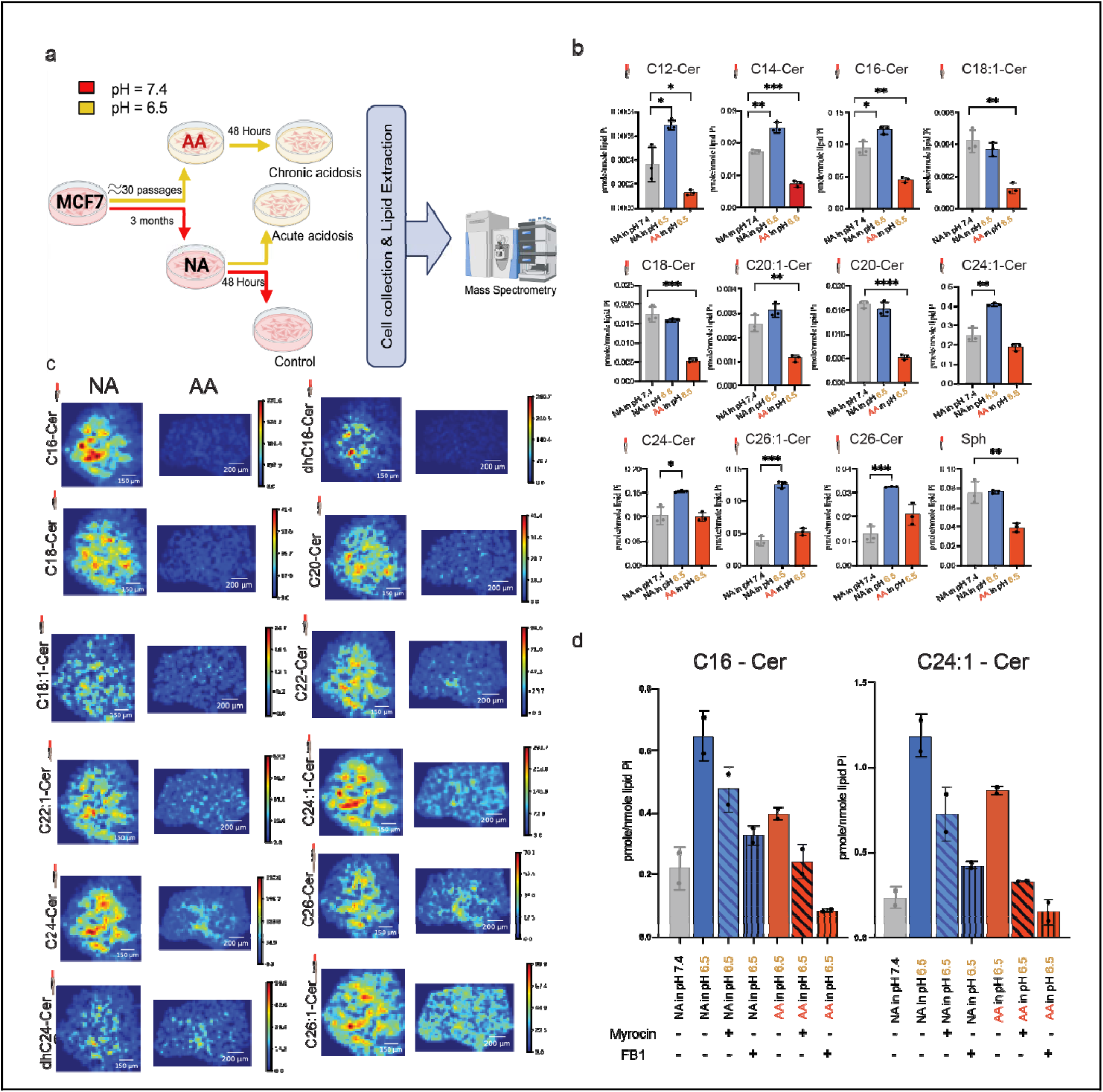
Chronic acidosis induces stable rewiring of ceramide metabolism and lipid spatial distribution. **a**, experimental design to measure SLs using mass spectrometry in acutely and chronically acid exposed MCF7 cells. Control cells were maintained at physiological pH (7.4) with the passage number matching the acid adapted cells. **b**, LC/MS of absolute values of Ceramides in MCF7 exposed to acid acutely and chronically compared to non-treated cells. **c**, MALDI imaging cross comparison of the same Ceramide in NA-MCF7 and AA-MCF7 spheroid. **d**, Mass spectrometry measurements of MCF7 cells in physiological pH and acid pH treated with Myriocin (100 nM, 48 hours) and FB1 (50 µM, 48 hours). Results are presented of the mean of two biological replicates with four technical replicates each. Statistical analysis was performed using a one-way ANOVA with multiple comparisons test. (*P < 0.05, **P < 0.01, ***P < 0.001, ****P < 0.0001, and ns p > 0.05).

We next performed MALDI-MS on 3D spheroids derived from NA-MCF7 and AA-MCF7 cells. Spheroids were embedded, sectioned, and mounted on the same slide to enable simultaneous analysis and direct comparison of the spatial distribution of identical ceramide species across adaptation states. This analysis enabled pixel-by-pixel cross-comparison of ceramide abundance within equivalent microenvironmental zones of the spheroids. In NA-MCF7 spheroids, multiple ceramide species were highly enriched in the inner spheroid regions corresponding to hypoxic and acidic zones, similar to Figure 2. In contrast, AA-MCF7 spheroids exhibited a marked reduction in ceramide signal across these same regions (Fig. 5c). Notably, the decrease was observed across nearly all detected ceramide species and was most pronounced in the spheroid core, where acidic stress is maximal. These spatially resolved findings closely mirrored the reductions observed in 2D lipidomic analyses, thereby independently validating the 2D MALDI and LC/MS results within a 3D tumor ecosystem context.

We next investigated whether ceramide production under acute and chronic acidosis is driven primarily by the de novo or salvage pathway. To address this, we used myriocin, an inhibitor of serine palmitoyltransferase that blocks de novo sphingolipid synthesis, and fumonisin B1 (FB1), which inhibits ceramide synthases and thereby disrupts ceramide generation through both de novo and salvage pathways. For both C16-Cer and C24:1-Cer, myriocin reduced ceramide levels under acute and chronic acidic conditions, indicating that de novo synthesis contributes to ceramide production (Fig. 5d, Extended Fig. 7). However, FB1 caused a substantially greater reduction in both species, suggesting that ceramide synthase-dependent salvage pathways also contribute strongly to the ceramide pool and that ceramide production in cells under acidosis is not sustained by de novo synthesis alone.

### Single-nuclei multi-omics reveals acidosis-dependent reprogramming of sphingolipid metabolic states

Our metabolic analyses indicated that acidosis induces extensive sphingolipid pathway rewiring and that key features of this phenotype are maintained during chronic adaptation. We therefore asked whether these metabolic changes are associated with transcriptional reprogramming through epigenetic regulation, and if so, whether such reprogramming represents a transient response to acute acid stress or a more stable adaptive state in chronic acidosis. To address this, we performed single-nuclei multi-omics (snRNA/ATAC-seq) of MCF7 cells under four environmental conditions: Control (pH 7.4), Acute exposure (NA MCF7 cells cultured in pH 6.5 for three days), AA-MCF7 (maintained at pH 6.5 for over three months), and Return (AA-MCF7 cells returned to physiological pH for three days). This design enabled us to distinguish transient stress responses from stable adaptive states and to assess whether acid-adapted cells retain memory of prior exposure. Notably, inclusion of the Return condition allowed direct interrogation of reversibility, providing a framework to determine whether the observed phenotypes reflect plastic responses or sustained adaptation.

A central challenge in analyzing sphingolipid pathway activity from single-nuclei data was the low and sparse expression of most sphingolipid metabolism genes. Many enzymes in this network, such as ceramidases, sphingomyelin synthases, and ceramide kinase, produce few detectable transcripts per cell, resulting in noisy and unreliable estimates of pathway activity at the single-cell level. To mitigate this technical limitation and stabilize gene-level signal estimates, we aggregated transcriptionally similar single nuclei into metacells using the hdWGCNA framework, pooling approximately 25 neighboring cells with similar expression profiles into single units (Morabito, et al. 2023; Methods).

UMAP visualization of the metacell transcriptomic profiles revealed clear separation between AA and and NA (Control) populations, whereas the short-term perturbations (Acute and Return) produced detectable but comparatively smaller shifts in transcriptional state (Fig. 6a). We then turned our attention into SLs metabolism related regulation. Ceramide, a central hub in this network, can be metabolized and removed through four major downstream pathways (Fig. 3c and 3h). These four metabolic directions can be conceptualized as “axes” of ceramide removal. While our inhibitor experiments demonstrated that all four routes contribute to ceramide clearance at the population level, whether individual cells prefer a single axis or simultaneously distribute metabolic flux across multiple axes remained unclear. To address this, we computed a “regulatory pressure” score, a measure of the net intensity with which upstream regulatory proteins collectively drive or suppress a given ceramide clearance axis, for each axis in each metacell. Projection of metacells colored by dominant axis assignment revealed that the majority of metacells align predominantly with a single sphingolipid axis, although a subset displayed ambiguous activity patterns consistent with simultaneous multi-axis engagement seen in inhibitor experiments (Fig. 6b). Quantification of the proportion of metacells assigned to each axis showed that the distribution of preferred metabolic directions differed substantially across conditions (Fig. 6c). As a proportion plot, this analysis reflects relative enrichment and shifts in population composition rather than absolute increases in pathway activity. Control cells showed relative enrichment of the C1P-associated axis, although this likely reflects a transcriptional state assignment rather than the dominant metabolic fate of ceramide. Acute acidosis shifted the population strongly toward the SM axis, while chronic acidosis preserved this SM-enriched pattern and further increased the fraction of GSL-aligned cells, consistent with a more polarized distribution of ceramide-utilization states during long-term adaptation. Notably, return exposure did not restore the control distribution, but instead was marked by expansion of the S1P-aligned population together with persistent GSL representation, suggesting that release from acidic stress produces a distinct transcriptional state rather than a full return to baseline. Our inhibitor experiments (Fig. 4) demonstrated that cancer cells dynamically reroute sphingolipid flux when individual clearance pathways are blocked, revealing substantial metabolic plasticity at the population level. To assess whether this plasticity is also reflected in the heterogeneity of metabolic states across individual cells, and whether acid adaptation alters the degree of such heterogeneity, we quantified metabolic diversity at two levels. At the single-metacell level, we computed the Shannon entropy of the four axis pressure scores, yielding a “metabolic diversity” index for each metacell. While the mean diversity was broadly similar across conditions, non-adapted populations (Control and Acute) exhibited substantially higher variance than AA populations (Chronic), indicating that chronic acid adaptation constrains cells toward a more consistent metabolic axis preference (Fig. 6d). At the population level, we computed the joint entropy across all four axes using quantile-binned pressure scores, which captures the overall phenotypic diversity of each condition’s cell population. Population-level diversity was higher in non-adapted cells (Control and Acute) compared to chronically adapted cells (Chronic) (Fig. 6e). Notably, both the Acute and Return perturbations were associated with transiently elevated diversity relative to their baseline states, suggesting that short-term environmental shifts temporarily expand the range of sphingolipid metabolic states that cells explore, consistent with the concept of metabolic plasticity as an immediate response to environmental perturbation. Finally, we sought to identify transcriptional regulators associated with each sphingolipid axis and whether these regulators differ between environmental conditions (Methods). This multi-evidence integration identified several candidate regulators associated with each axis (Fig. 6f-j), including transcription factors with condition-dependent activity patterns. No single dominant master regulator emerged for any axis; instead, multiple transcription factors displayed moderate and context-dependent associations. The axis-specific and condition-dependent transcription factor signatures observed here, including enrichment of FOX, ZNF, and MEF family members, are consistent with regulation through chromatin-dependent transcriptional programs. Such an architecture would enable rapid and reversible switching between metabolic states without requiring stable genetic changes, providing a mechanistic basis for the observed metabolic plasticity. These results suggest that the transcriptional regulation of sphingolipid metabolic states is governed by a distributed, combinatorial regulatory architecture rather than by a single central switch supporting our hypothesis as metabolic flux governing the whole activity as of cancer cells metabolic plasticity.

**Figure 6.**
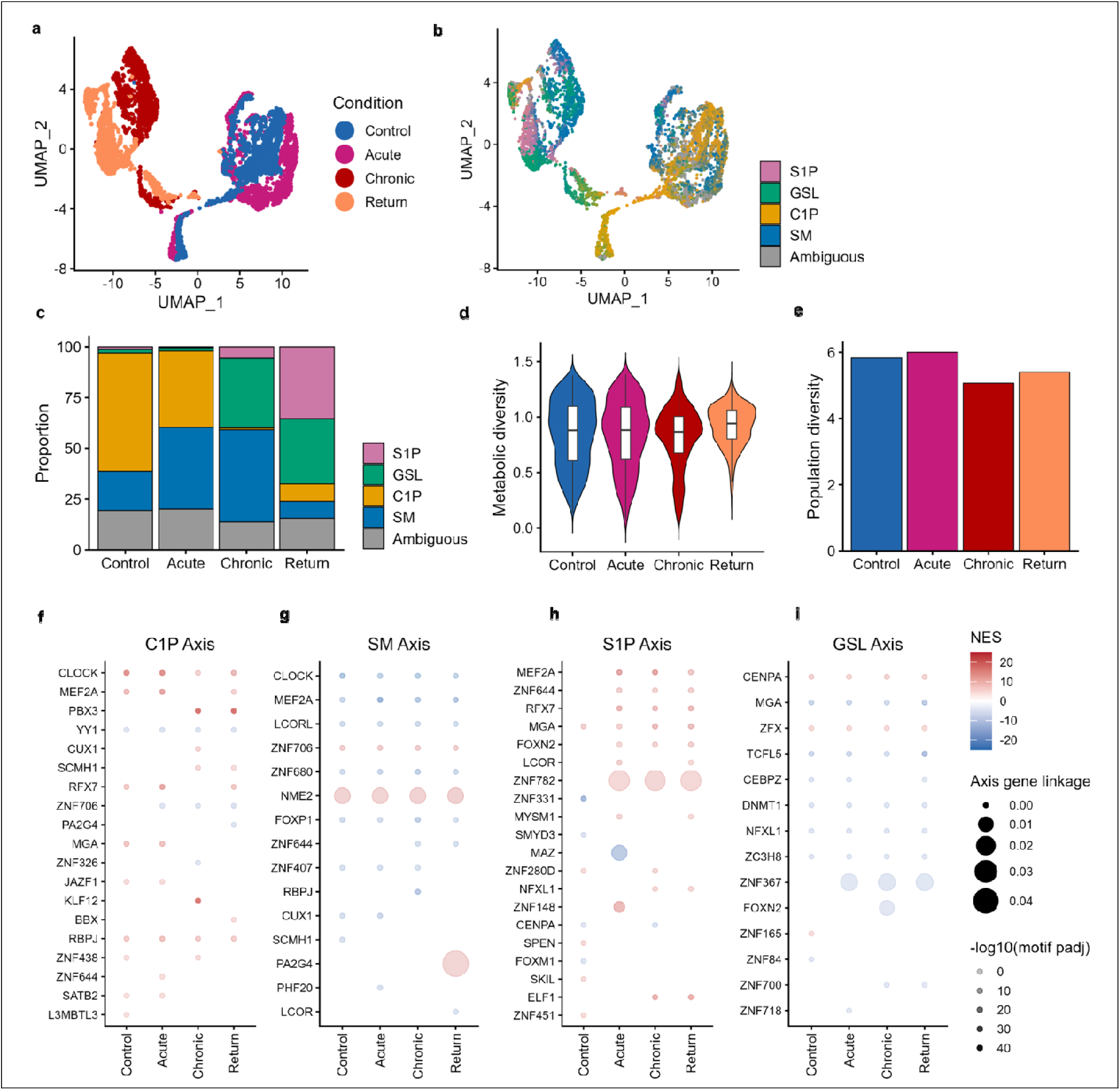
Single-nuclei multi-omics analysis identifies condition-specific sphingolipid axis programs across acute, chronic, and return states. **a**, UMAP projection of single-cell transcriptional states from MCF7 cells under Control, Acute acidosis, Chronic acidosis, and Return conditions, revealing distinct but partially overlapping cell populations. **b**, Conceptual model of sphingolipid metabolic plasticity centered on ceramide (Cer), illustrating dynamic redistribution of metabolic fl x toward alternative pathways-S1P, GSLs, SM, and C1P - under changing acidic environmental conditions. Bidirectional arrows indicate reversible transitions between metabolic states. **c**, UMAP colored by dominant sphingolipid metabolic axis assignment, highlighting favorite metabolic phenotypes under four different conditions of acid treatment, including S1P-, GSL-, C1P-, SM-dominant, and ambiguous states. **d**, Proportional distribution of metabolic states across conditions, showing shifts in pathway utilization during acute and chronic acidosis and partial reversion upon return to physiological conditions. **e**, Violin plots of metabolic diversity across conditions, indicating changes in intra-population metabolic heterogeneity. **f**, Population diversity scores across conditions, demonstrating reduced diversity under chronic acidosis and recovery upon return. **g–j**, Transcription factor motif enrichment associated with each sphingolipid metabolic axis (C1P, SM, S1P, and GSL), showing condition-specific regulatory programs. Dot size represents motif significance (−log10 adjusted P value), and color indicates normalized enrichment score (NES).

## Discussion

Tumor evolution unfolds within spatially structured ecosystems in which persistent microenvironmental stressors impose strong and predictable selective pressures(*30*). Among these, extracellular acidosis is a defining feature of many solid tumors, arising early during tumorigenesis and persisting throughout progression(*31–34*). Survival within acidic niches therefore requires adaptive strategies that operate on ecological timescales, enabling cancer cells to maintain fitness despite chronic stress and spatial heterogeneity(*35*, *36*). Our findings identify sphingolipid metabolism, specifically the regulation of ceramide turnover, as a central eco-evolutionary trait that enables cancer cell persistence in acidic tumor ecosystems.

A key insight from this work is that cancer cells do not adapt to acidosis by committing to a single optimal metabolic pathway. Instead, they exploit metabolic degeneracy, whereby multiple, biochemically distinct ceramide clearance routes converge on a shared functional outcome: avoidance of ceramide-induced cytotoxicity and preservation of growth(*37*). Degeneracy is a hallmark of robust adaptive systems and is widely observed in biological networks that must remain functional under fluctuating or unpredictable conditions(*38*). In the tumor context, this property provides resilience against both environmental stress and targeted perturbations, allowing cancer cells to maintain fitness even when individual metabolic routes are disrupted(*39*).

Our temporal analyses further reveal that adaptation to acidosis is not static. Acute acid exposure induces ceramide accumulation, consistent with ceramide’s established role as a stress-responsive, pro-apoptotic lipid that reduces cellular fitness(*40*, *41*). Chronic acidosis, however, selects for phenotypes capable of mitigating this liability through sustained metabolic rewiring, resulting in reduced ceramide levels and increased flux through alternative sphingolipid pathways. This transition reflects an eco-evolutionary process in which an initially deleterious phenotype exposes variation upon which selection can act, favoring plastic responses that stabilize fitness under long-term stress. Importantly, the adaptive advantage conferred by sphingolipid plasticity appears to lie not in maximizing short-term growth, but in minimizing the risk of catastrophic failure(*42*, *43*). This aligns with principles of bet-hedging, in which organisms adopt strategies that reduce variance in fitness across time or space, even at the expense of peak performance(*44*, *45*). By maintaining access to multiple ceramide clearance routes, cancer cells effectively insure themselves against sudden environmental shifts or pathway-specific perturbations. Our observation that single-pathway inhibition rarely compromises viability, whereas simultaneous closure of all clearance routes is lethal, is consistent with selection favoring strategies that maximize long-term, or geometric mean, fitness in harsh and heterogeneous environments(*45*, *46*). Spatial analyses further emphasize that this adaptive strategy is context dependent. Distinct sphingolipid phenotypes emerge across tumor habitats, constrained by local resource availability and stress intensity. Such spatial partitioning promotes the coexistence of multiple metabolic states within the same tumor, increasing phenotypic diversity and enhancing evolvability. This spatially maintained diversity provides a reservoir of adaptive potential that can be drawn upon during progression or therapy, offering a mechanistic explanation for the resilience of acidic tumors to metabolic intervention.

From a therapeutic perspective, these findings underscore a fundamental limitation of pathway-centric targeting strategies. In degenerate, plastic systems, inhibiting a single metabolic route is unlikely to produce durable benefit, as cells can rapidly reroute flux through alternative pathways. Instead, effective intervention may require strategies that constrain adaptive flexibility itself, by reducing network connectivity, increasing the cost of metabolic switching, or inducing evolutionary traps that exploit the liabilities of plastic states. Targeting metabolic plasticity, rather than individual pathways, may therefore represent a more effective approach to limiting tumor adaptation and evolutionary persistence(*47*). Single-cell transcriptomic analysis identified multiple transcriptional regulators associated with distinct sphingolipid metabolic states, including members of the FOX, ZNF, and MEF families, suggesting that metabolic plasticity is accompanied by coordinated, state-dependent transcriptional programs. However, the axis-specific and condition-dependent nature of these signatures, together with the absence of a single dominant regulator, indicates that control of sphingolipid flux is unlikely to be governed by a linear transcriptional hierarchy. Instead, these data are consistent with a model in which metabolic state transitions are enabled by dynamic changes in chromatin accessibility that permit selective engagement of alternative transcriptional modules(*48*). Such an architecture would allow rapid and reversible switching between ceramide clearance routes without requiring stable genetic alterations, thereby supporting the observed plasticity under acidic selection. While our current analyses infer transcription factor activity, they point to chromatin-level regulation as a key layer of control. Defining the epigenomic landscape underlying these transitions—through direct measurements of chromatin accessibility and regulatory element usage—represents an important next step toward understanding how metabolic plasticity is established and maintained in tumor ecosystems.

### Model: sphingolipid plasticity as an adaptive fitness landscape

We propose a model in which ceramide occupies a central position within an adaptive metabolic landscape shaped by acidic selection. The four major ceramide clearance routes define alternative adaptive phenotypes whose fitness values are similar in magnitude but highly interconnected. Chronic acidosis selects not only for movement away from ceramide-rich states but also for increased landscape navigability, enabling rapid transitions between routes when perturbations occur. In this landscape, fitness is maintained through redundancy and connectivity rather than optimization of a single pathway. Therapeutically, this predicts that restricting landscape connectivity, simply by collapsing multiple escape routes or targeting the mechanisms that enable switching, may force cancer cells into low-fitness valleys, reducing adaptation and limiting evolutionary resilience.

## Materials and Methods

### Cell Culture and acid adaptation

MCF-7 cells were maintained under standard culture conditions (37°C, 5% CO) in RPMI (Corning) supplemented with 5% FBS (Corning). Growth medium was further supplemented with 12.5 mmol PIPES, 12.5 mmol HEPES and the pH adjusted to 6.5. Cells were tested for mycoplasma contamination monthly and authenticated using short tandem repeat (STR) DNA profiling, following ATCC’s guidelines. To achieve acid adaptation cells were chronically cultured and passaged upon reaching 80% confluency directly in pH 6.5 medium for approximately 3 months. Control cells or (non-adapted cells) were cultured in parallel in pH 7.4 medium to obtain a control group with the same passage number as AA cells. Acid-adapted (AA) and non-adapted (NA) cells underwent about 35 passages.

For inhibitor studies, the following compounds were used: Fumonisin B1 (50 µM, BML-SL220) as a ceramide synthase inhibitor, Myriocin (100 nM, M1177, Sigma) as an SPT inhibitor, PF-543 (100 nM, 567741, Calbiochem) as an SK1 inhibitor, Eliglustat (200 nM, M15398) as a glucosylceramide synthase inhibitor, HPA-12 (10 µM, 28350) as a CERT inhibitor, and NVP-231 (500 nM, 13858) as a ceramide kinase-1 inhibitor.

### 3D Spheroids processing

We grew spheroids using U-bottom 96 well plates with at least 10 replicates per condition. 2000 cells per well were seeded in a sterile, ultra-low-attachment U-bottom 96-well plate and incubated under standard culture conditions (37°C, 5% CO). The following day, the plate was centrifuged at 18 x g for 5 minutes at 25°C to initiate cell aggregation within each well. The plate was then incubated for an additional 2 days. After this incubation period, the medium was changed, and 5% Matrigel (356237, Corning) was added to each well. The plate was centrifuged again at 18 x g for 5 minutes at 25°C. The plates were imaged every 24 hours using Cytation 10 automated imaging, and the sectional area of the spheroids was measured to generate growth rate curves using (Software “Gen5”). On the final day, the spheroids were collected, washed in PBS, embedded in a gelatin mold and instantly frozen. Cryostat is used to cut the blocks into 4um sections for HE staining, followed by 4um sections for MACSima mIF staining, and finally 10 um sections for MALDI in negative and positive mode for both small and big molecules ranging from 100 to 1600 m/z.

### Mass spectrometry-based lipids measurement in cancer cells

Cells were collected by centrifugation at 800*g* for 5 min. Then, cells were washed one time in cold PBS and lysed in 2 mL of cell extraction mix (2:3 70% Isopropanol:Ethanol). To perform sphingolipid quantification, samples were spiked with the corresponding internal standards (50 pmol), and extracts were then analyzed by the Lipidomic Core Facility at Stony Brook University Medical Center, as described previously. Sphingolipid species were identified on a Thermo Finnigan TSQ700 triple quadrupole mass spectrometer. Sphingolipids from cellular extracts were normalized to total lipid phosphates present in the cells after a Bligh and Dyer extraction.

Spheroids were washed with PBS and fixed in 5% PFA for 30 minutes at room temperature. They were then embedded in 10% Gelatin using a plastic mold and stored at -80 °C. Cryo-sectioning was then performed on a CM1950 cryostat (Leica Biosystems GmbH, Nussloch, Germany).

Frozen blocks were mounted with carboxymethyl cellulose (CMC) on the sample holder and were kept inside the cryostat chamber (temperature set to -20 °C) for 15 mins prior to sectioning. For H&E staining, 5 µm thick sections were used right after sectioning. For MALDI-MSI analysis, spheroid sections at 10 µm thickness were cut, and were mounted on IntelliSlides (Bruker Daltonics, Bremen, Germany) while being kept frozen. During subsequent sample preparation, frozen slides were dried in a vacuum desiccator for ∼30 mins. Three matrices were applied on sequential spheroid sections, including 9-Aminoacridine (9AA) (5 mg/ml in 70 % Ethanol (EtOH)) for analyzing metabolites, 1,5-Diaminonaphthalene (DAN) (5 mg/ml in 90 % Acetone) for ceramides, and 2,5-Dihydroxybenzoic acid (DHB) (40 mg/ml in 70 % Methanol (MeOH) and 0.1% Trifluoroacetic acid (TFA)) for sphingomyelins. The matrices were sprayed using an HTX-TM5 sprayer following the methods provided by HTX Technologies, LLC, Chapel Hill, North Carolina, United States. The matrices were sprayed using an HTX-TM5 sprayer following the methods provided by HTX Technologies, LLC, Chaepl Hill, North Carolina, United States. Key spraying parameters for matrix application are summarized in the table below:

Embedding and cryo-sectioning of spheroids and sample preparation for MALDI-MSI.

| Matrix | Temp (°C) | # Passe (Count) | Conc (mg/ml) | Flow rate (ml/min) | Velocity (mm/min) |
| --- | --- | --- | --- | --- | --- |
| 9-AA | 90 | 8 | 5 | 0.12 | 1200 |
| DAN | 30 | 8 | 5 | 0.1 | 1200 |
| DHB | 75 | 8 | 40 | 0.1 | 1200 |

### MALDI mass spectrometry imaging

MALDI-MSI experiments were performed on a timsTOFfleX mass spectrometer (Bruker Daltonik, Bremen, Germany), operated in negative ion mode for detection of ceramides/metabolites and positive ion mode for sphingomyelins. MALDI-MSI data was acquired within the mass-to-charge ratio range of 50-2000 m/z with a spatial resolution of ∼20 μm, and with the optimized experimental settings previously reported in reference (*49*), where validation of MADLI signals for lipids were conducted on the same instrument used in this study. All the raw data was processed in SCiLS Lab (Version 2023a Pro) to generate ion images and mass spectra, which were normalized to the total ion count. The corresponding imzML and ibd files extracted from SCiLS Lab were imported into METASPACE and were processed for non-targeted analysis.

### Targeted MALDI analysis

For the MALDI image analysis, the coregistration of the three sequential cuts for sphingomyelin (SM), ceramides (Cer), and metabolites was performed using affine transformation. This process involved aligning the bounding boxes of each cut to ensure accurate spatial correspondence across the different molecular images. Following the alignment, the colocalization score for each pair of molecules was calculated using Pearson’s correlation coefficient. Pairs exhibiting a correlation score above the threshold of 0.7 were considered significant and visualized accordingly. Additionally, analysis was conducted to compare the intensity distribution of pixels in the outer layer of spheroids, defined as within 150 μms from the boundary, with that of the inner layer. A Wilcoxon rank-sum test was applied to determine if there were significant differences between the intensity distributions of the outer and inner layers. Code for the image analysis is available at: https://github.com/sbu-damaghi-ceel/Lipid_analysis.

### Non-targeted MALDI Analysis

A cloud-based platform, METASPACE was utilized for the initial processing of MALDI mass spectrometry imaging data. METASPACE performs peak picking, filtering, and annotation. Specifically, the platform imported raw MALDI imaging datasets, measured spatial chaos, spatial isotope and spectral isotope; calculated a False Discovery Rate (FDR) for each m/z annotation; enabled selection of reliable m/z features by applying a user-defined FDR threshold (10% in this analysis). The resulting high-confidence m/z list served as the basis for downstream clustering and phenotypic analysis. And the uploaded dataset is publicly accessible at https://metaspace2020.eu/project/88020884-c6d1-11ef-a046-97350f84f65a?tab=datasets.

### SPHERE Plot

To compare habitats across all experimental conditions, MALDI metabolomic and lipidomic intensities were normalized relative to the normoxic habitat of the control samples treated with DMSO, where normoxic habitats were defined as 125 μms from the boundary and hypoxic habitats were defined as >125 μms from the boundary. A heatmap of the normalized intensities was plotted on a circular axis, with radial sections corresponding to distinct habitats (normoxic and hypoxic) and angular divisions representing different molecules grouped by their biochemical class (metabolites, ceramides, and sphingolipids). Each condition (e.g., DMSO control, PF, EL, HPA, NVP) was visualized in separate plots to highlight changes in spatial intensity distributions. Statistical significance between the normoxic habitat of the control samples and the habitats across all conditions was assessed using the Mann-Whitney U test. To further illustrate molecular responses, bars overlaying the heatmap depict fold changes relative to the control normoxic habitat, with bar height representing the magnitude of the fold change and color indicating directionality (red for positive, blue for negative) and statistical significance (gray for nonsignificant, *p* > 0.05).

### Construction of Sphingolipid CRISPR Library

For dropout screening studies of the sphingolipid network, a pooled library of gRNAs targeting 39 core enzymes of the sphingolipid network was developed (see Supplementary table S2 for list of genes). For this, six gRNA sequences per gene as well as 30 non-targeting control gRNA sequences were obtained from the GeCKO library. These sequences were complemented by an additional 4 independent sequences per gene designed using the CHOPCHOP algorithm. Sequences were synthesized as pooled oligos and subsequently amplified and subcloned into Lenticrispr v2.0 by Gibson cloning according to the methods of Sanjana et al. (*50*). The ligation product was transformed into 25u of electrocompetent cells, with 10 parallel reactions performed to ensure no loss of representation. Following transformation, E. coli were plated onto 245 mm x 245 mm plates (Corning) with carbenicillin selection (50 ug/ml). The next day, all colonies were scraped and combined for plasmid DNA extraction using a endotoxin free maxiprep kit from Qiagen. For validation of the library, purified plasmid was subjected to nested PCR to amplify inserts, and a second round of PCR to add sequencing primers. PCR products from four independent reactions were purified, pooled, and sequenced by Novogene. Analysis of sequences by Bowtie (allowing for up to 1bp mismatch) showed presence of 99.2% of designed gRNAs with 85% of gRNAs in a 15-fold range (from 2000 to 30000 reads).

### Sphingolipid Lentiviral Generation

To generate lentiviral particles, Sphingolipid gRNA Library was co-transfected with 1.5 μg or each viral packaging plasmid (VSV-G and dVPR) in 293T cells (2000k cells per 10cm dish) using Lipofectamine 2000 according to the manufacturer’s protocol. Cells were incubated overnight with the virus, media was changed after 24 hrs, and 24 hrs later 4 mL additional media was added. At 72 hours following transfection, viral media was collected, filtered (Millipore 0.45 μm PVDF), and aliquoted for storage at - 80C until use.

### Synthetic Lethality Screen

For the negative selection screen, 2 x 10^5^ MCF7 cells were infected with Sphingolipid lentiviral supernatant to ensure MOI of ≤ 0.3 and coverage of ≥ 300 cells expressing each gRNA. Cells were selected with puromycin (1.75 μg/mL for 10 days) to ensure viral integration and depletion of gene products. Following antibiotic selection, one aliquot of (500k) cells was collected (as a day 0 control sample) and remaining cells (500k) were placed in either the normal or acidic environment. After 48 hrs, cells were collected and all cell aliquots were harvested for gDNA via Qiagen QIAamp DNA Micro Kit (56303). Following DNA extraction, the gRNA cassettes were amplified via PCR (Supplementary Table S2). After pooling like samples, Illumina indexing primers were added via additional PCR (Supplementary Table S2) and PCR product was purified via Qiagen Qiaquick PCR Purification Kit (28104). Purified DNA was quantified via Qubit and equimolar amounts were pooled and submitted to Novogene for sequencing (NovaSeq X Plus Series).

### Sequencing and Statistical Analysis

To assess the enrichment or depletion of each gRNA and normalize data, MAGeCK-MLE (version 0.5.9.5) statistical package and MAGeCKFlute (version 2.9) were utilized respectively. Briefly, read counts of each sample were compared to the read count from the control to generate beta scores. Differential beta scores were calculated by assessing the difference between the two experimental conditions. Experiments were conducted in biological quadruplet.

### Real time PCR

Total RNA from cells was extracted using an Rneasy extraction kit (Qiagen) in accordance with the manufacturer’s protocol. cDNA was synthesized from 1.0 µg of RNA using Applied Biosystems High-Capacity cDNA Reverse Transcription Kit (43-688-14) according to the manufacturer’s instructions. Real-time PCR analysis was performed on a QuantaStudio 7 Flex Real-Time PCR system using Applied Biosystems PowerTrack SYBR Green Master Mix (Cat# A46012). Expression levels of mRNA were measured as a ratio with β-actin, which was the normalization control.

**Table 1.**
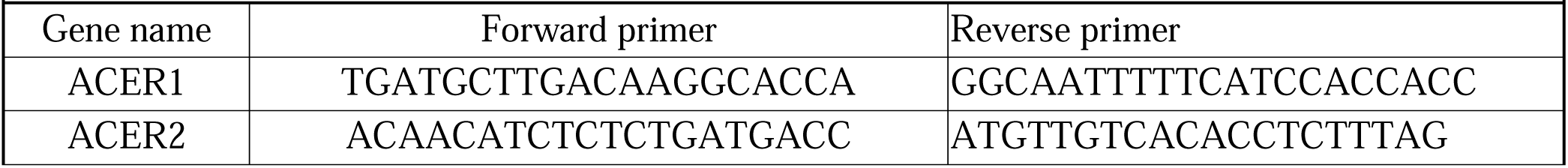
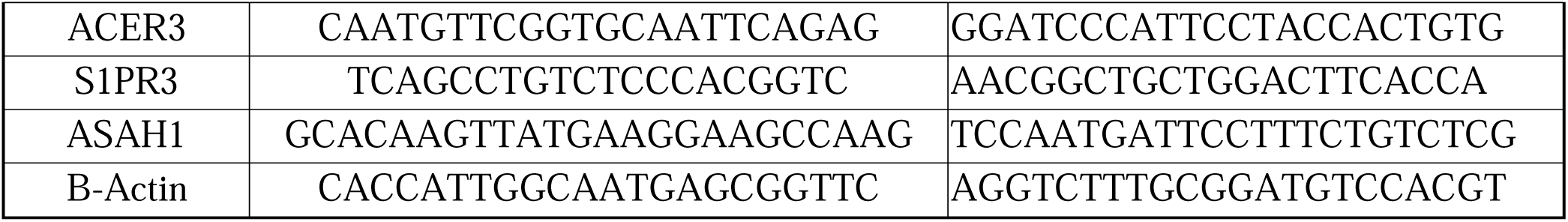
Primer sequences.

### Immunofluorescence

NA-MCF7 and AA-MCF7 cancer cells were seeded in 8 well chamber slides and treated with media at pH 7.4 or pH 6.5 for 48 hours. Then media was removed and cells were rinsed with PBS, fixed in 4% paraformaldehyde for 10 mins at room temperature, and then blocked with 4% bovine serum albumin in PBS. Samples were incubated with SK1 rabbit polyclonal primary antibody (1:1,000, PIPA522994, Invitrogen) and secondary Alexa-Fluor 488 goat anti-rabbit (1:250) antibody. Coverslips were mounted using Flurosheild Mounting Medium (Abcam, Cat# NC0200574). Images were captured using Cytation10 microscope in confocal mode.

### Multiplex immunofluorescence (mIF) with the MACSima imaging platform

#### Sample preparation

Frozen embedded spheroids were cryosectioned with a CM3050 cryostat (Leica), 8 µm sections were mounted on SuperFrost Plus slides and stored in -80°c. On the day of use, the MACSwell™ Imaging frame was directly mounted onto the slide and immediately fixed with the appropriate volume of 4% PFA solution at room temperature for 10 minutes. The slide was then washed three times with MACSima Running Buffer.

For nuclear proteins, a membrane permeabilizing protocol was performed on the sample for better quality antibody binding and imaging. The sample was fixed in cold acetone for 5 minutes at room temperature. Following this, one wash with 0.1% Triton™ X-100 in PBS as well as 3-5 additional washes with MACSima Running Buffer was performed.

Immediately before running the sample in the MACSima Imaging Platform, the sample was stained with a 1:10 dilution of DAPI staining solution in the dark at room temperature for 10 minutes. Three additional washes with MACSima Running Buffer were performed, and the appropriate volume of buffer was added in accordance with the manufacturer’s protocol.

Antibodies list: Antibodies were diluted in MACSima Running Buffer in accordance with the manufacturer’s recommendations and/or after being optimized for the MACSima Imaging Platform. The following antibodies were used: Cytokeratin-APC (130-123-091), CAIX-AlexaFluor488 (sc-365900), ErbB2-PE (130-127-979), GLUT1-APC (130-127-114), GLUT3-FITC (sc-74399), CD107b-PE (130-118-818), beta-Catenin-APC (130-124-444), MCT4-AlexaFluor488 (sc-376140), CD324-PE (130-127-980), Vimentin-APC (130-127-018), Fibronectin-FITC (130-122-864), SPK1-FITC (NBP2-74332F), Progesterone Receptor-AlexaFluor488 (35591), Hif2a-PE (NB100-132AF594), Cleaved-PARP1-PE (130-128-159), Ki67-PE (130-119-356).

### MACSima Segmentation and Normalization

Cell segmentation was performed in MACSiQ_View software with the setting of ‘constrained donut’ mode which uses DAPI channel for nucleus and Pan CytoKeratine (PCK) channel as cytoplasmic marker to delineate the cell boundaries. Following segmentation, cell expression tables were exported, containing quantitative marker intensities for each segmented cell. These expression profiles underwent normalization on a per-slide, channel-specific basis, scaling intensities relative to the 95th percentile to mitigate slide-to-slide variability. Subsequently, a phenotyping pipeline— adapted from the Pixie(*51*) — was applied. This involved overclustering through a Self-Organizing Map (SOM) followed by consensus hierarchical clustering on the standardized (z-scored) cluster scores. The output clusters were then further refined and annotated using a graphical user interface. This systematic approach ensured that each cell was accurately and robustly phenotyped based on its multiplexed marker expression profile, facilitating robust downstream analyses of cellular heterogeneity.

### Phenotyping: MACSima and MALDI

A two-step clustering module, adapted from the Pixie framework (*51*) was employed for both MALDI and MACSima data analysis. This module involves an initial overclustering stage using a Self-Organizing Map (SOM) followed by a consensus hierarchical clustering step on the standardized (z-scored) cluster scores. The two-step clustering approach serves as a robust foundation for subsequent phenotyping efforts across different datasets.

<u>For MACSima phenotyping</u>, all control spheroids were analyzed. The workflow utilized the normalized cell expression tables derived from the segmentation process, as described earlier. The acid/hypoxia-specific panel, composed of markers CAIX, CAXII, GLUT1, GLUT3, LAMP2b, MCT4, HIF2a, Cleaved PARP and Ki-47, was used to characterize the phenotypes. The two-step clustering module generated preliminary cell clusters, which were further refined and annotated using a graphical user interface within MACSima. Manual refinement was pivotal at this stage, enabling experts to incorporate biological insights and ensure accurate phenotyping based on the multiplexed marker expression profiles.

<u>For MALDI phenotyping</u>, all pixels from the control spheroids were included, excluding small spheroids with a radius <125 micrometers. The molecule list was thresholded using METASPACE with an FDR ≤ 10%. A robust shared molecule features list, comprising 52 molecules common to all spheroids, was identified for phenotyping (Supplementary table S1). The same two-step clustering module was applied to this dataset, but manual refinement was kept to a minimum to maintain the analysis’ objectivity and reproducibility. This data-driven approach ensured that the phenotyping remained unbiased while focusing on metabolic subtype identification.

### Integration of MALDI and MACSima

The MALDI imaging and MACSima imaging were done on two sequential cuts of spheroids, to unify them into the same geometric space a manual co-registration step was done with FIJI BigDataViewer(*52*) extension. Once co-registered, each cell not only retained its MACSima-derived proteomic phenotype but was also assigned a corresponding MALDI-derived metabolomic phenotype based on its spatial overlap with defined metabolic regions. And a co-occurrence table could be computed linking MALDI-derived metabolic regions with MACSima-defined cellular phenotypes. Each entry in the co-occurrence table represents the frequency of cells with a specific proteomic phenotype residing within a particular metabolic region. This table provides insight into spatial associations between cellular functions and local metabolic environments.

### Patient derived organoids culture

Primary human tissue samples were collected with informed consent from all participants, adhering to ethical and institutional guidelines. Organoid collection and processing were performed according to the protocol outlined in Sachs et al. (*53*). Tissue fragments were minced into small pieces and transferred to a 10 mL conical tube containing digestion medium. Tissue digestion was performed using adDMEM/F12+++ medium supplemented with 5 mg/mL collagenase (Cat# C213G Sigma), with samples incubated on an incubated tube rotator at 37°C for 30–60 minutes. The digestion process was monitored under a microscope, with small organoid structures visible by the end of the incubation period. Following digestion, tissue suspensions were washed with adDMEM/F12+++ medium and centrifuged at 400 × g for 10 minutes at room temperature to pellet the cells. For samples containing a high amount of blood, the pellet was treated with red blood cell lysis buffer, incubated on ice for 5 minutes, and washed again with adDMEM/F12+++ medium to ensure purity of the epithelial components. After RBC lysis, samples were centrifuged at 50 × g for 1 minute to pellet organoids specifically, allowing separation from any remaining debris.

The resulting cell pellet was resuspended in Matrigel (356237, Corning) and plated as 20 µL droplets onto a pre-warmed 48-well plate. Plates were incubated at 37°C for 30 minutes to allow Matrigel solidification, after which 500 µL of pre-warmed organoid medium, supplemented as detailed in the Supplementary Table S3, was added to each well. The culture medium was refreshed every 2–3 days, and the plates were imaged regularly to monitor organoid formation and growth.

### Single-nucleus multi-omics data generation and preprocessing

MCF7 cells cultured under four conditions (Control at pH 7.4; Acute: non-adapted cells exposed to pH 6.5 for 3 days; Chronic: maintained at pH 6.5 for >3 months; Return: acid-adapted cells returned to pH 7.4 for 3 days) were profiled using the 10x Genomics Multiome ATAC + Gene Expression platform. Libraries were prepared according to the manufacturer’s protocol and sequenced on an Illumina NovaSeq instrument. Raw sequencing data were aligned to the GRCh38 (hg38) human reference genome and processed using Cell Ranger ARC (10x Genomics) to generate per-sample filtered feature-barcode matrices and ATAC fragment files.

### Seurat object construction and quality control

For each sample, the filtered feature-barcode HDF5 file was read using Seurat v5 ([3] Hao et al., 2024), and separate RNA and ChromatinAssay ([4] Signac; Stuart et al., 2021) objects were created from the Gene Expression and Peaks modalities, respectively. Gene annotations were derived from EnsDb.Hsapiens.v86, with sequence levels standardized to UCSC notation (hg38). Per-nucleus quality control metrics were computed, including nucleosome signal and TSS enrichment (Signac) for the ATAC modality, and mitochondrial gene percentage for the RNA modality. Nuclei failing any of the following thresholds were removed: ATAC fragment count between 1,000 and 100,000; nucleosome signal < 2; TSS enrichment > 1; RNA UMI count between 1,000 and 35,000; mitochondrial gene fraction < 20%. After filtering, all mitochondrial genes were removed from the RNA count matrix to prevent confounding of downstream expression analyses.

### Peak calling and unified peak set construction

Accessible chromatin peaks were called independently for each sample using MACS3 ([5] Zhang et al., 2008) via the Signac CallPeaks interface applied to the ATAC assay. Called peaks were filtered to retain only those on standard chromosomes and to exclude hg38 unified blacklist regions ([6] Amemiya et al., 2019). A union peak set was generated by merging peaks across all four samples using GenomicRanges::reduce, and a unified peak-by-cell count matrix was constructed for each sample by quantifying fragment overlap with the union peak set using Signac::FeatureMatrix. The resulting unified ChromatinAssay (named “peaks”) was appended to each Seurat object alongside the original Cell Ranger–derived ATAC assay.

### RNA and ATAC transformation and multimodal integration

All four per-sample objects and the merged object were processed through parallel transformation pipelines. For the RNA modality, expression data were normalized using (*54*), followed by PCA (50 components), shared nearest neighbor (SNN) graph construction, and UMAP embedding. For the ATAC modality, the unified peaks assay was processed using term frequency–inverse document frequency (TF-IDF) normalization, followed by singular value decomposition (SVD; 51 components with the first component excluded, as is standard practice, to remove the technical correlation with sequencing depth) and UMAP embedding. The RNA and ATAC modalities were then integrated using Seurat’s weighted nearest neighbor (WNN) framework, which constructs a joint cell neighborhood graph from the SCT-derived PCA and LSI embeddings with learned per-cell modality weights, followed by multimodal UMAP embedding.

### Clustering with automated parameter optimization

Clustering was performed on both the SCT-derived PCA and the peaks-derived LSI embeddings using a grid search procedure inspired by the ACDC framework(*55*). For each modality, Leiden clustering (algorithm 4 in Seurat) was applied across a two-dimensional grid of k-nearest-neighbor values (k = 11, 21, 31, …, 101) and resolution parameters (resolution = 0.1, 0.3, 0.5, …, 1.9). For each parameter combination, silhouette scores were computed on the corresponding Euclidean distance matrix (PCA or LSI embedding space), and the mean of per-cluster mean silhouette widths (ACDC-style unweighted average) was used as the optimization criterion. The parameter combination yielding the highest silhouette score was selected, and final cluster assignments were stored as acdc_sct (RNA-based) and acdc_peaks (ATAC-based) in the object metadata.

### Transcription factor motif accessibility analysis (chromVAR)

Transcription factor motif accessibility deviations were estimated across all nuclei using chromVAR (*56*). Position frequency matrices (PFMs) for human transcription factors were obtained from the JASPAR 2024 CORE vertebrate collection(*57*), restricted to Homo sapiens (taxon ID 9606). Motif occurrences within the unified peak set were identified using the hg38 reference genome (BSgenome.Hsapiens.UCSC.hg38), and per-nucleus deviation z-scores were computed and stored as a separate chromVAR assay.

### Metacell construction

Because sphingolipid metabolism genes are generally expressed at low levels in single-nucleus RNA-seq data, with many key enzymes detected only sporadically across individual nuclei, direct estimation of pathway activity from raw single-cell profiles is unreliable. To overcome this signal sparsity, transcriptionally similar nuclei were aggregated into metacells using the hdWGCNA R package (*58*). Metacells were constructed by summing RNA raw counts across k = 25 nearest neighbors in the SCT-derived PCA space, with a maximum of 10 shared cells permitted between metacells and a minimum group size of 30 cells. This aggregation procedure averages out stochastic dropout and amplifies coherent transcriptional signals while preserving biological heterogeneity across cell states and conditions. The resulting metacell expression profiles were then used as the basis for all downstream regulatory network inference and pathway activity estimation. Grouping was stratified by both experimental condition and SCT-derived cluster identity (acdc_sct) to preserve biological heterogeneity across cell states. The resulting metacell count matrix was normalized to counts per million (CPM) for downstream analyses. Metacell-level chromVAR deviation scores were computed by averaging the per-nucleus chromVAR z-scores across all constituent nuclei of each metacell using the metacell membership assignments stored by hdWGCNA.

### Gene regulatory network inference and transcription factor activity estimation

For each transcription factor in the ARACNe3-inferred regulatory network, we first estimated its protein-level activity from the expression patterns of its downstream target genes using the VIPER algorithm (*59*), which infers how actively a given regulator is driving its targets in each metacell based on the coordinated expression changes across its regulon. The regulatory pressure score then integrates these inferred activities across all transcription factors that regulate a given axis’s gene set, weighted by their regulatory strength (mode of regulation and likelihood), thereby estimating the net transcriptional drive toward each ceramide clearance route. A transcriptional regulatory network was inferred from the metacell CPM expression matrix using ARACNe3 (*60*, *61*)https://github.com/califano-lab/ARACNe3). The regulator set consisted of human transcription factors curated from the Human TFs database (*62*); humantfs.ccbr.utoronto.ca, v1.01), filtered to those present in the metacell expression matrix. ARACNe3 was run with a significance threshold of α = 0.05, 30 bootstrap iterations, 8 threads, and a fixed random seed of 1234. The consolidated network was reformatted to a three-column format (regulator, target, mutual information) for compatibility with the VIPER R package (*59*). A regulon object was generated using the aracne2regulon function, and VIPER was applied to the metacell CPM matrix using the rank-based method with a minimum regulon size of 25, producing a normalized enrichment score (NES) for each transcription factor in each metacell. The VIPER activity matrix was stored as a separate assay within the metacell Seurat object. Independent dimensionality reduction (PCA, 30 components; UMAP) was performed on both the SCT-normalized metacell expression data and the scaled VIPER activity matrix.

### Sphingolipid axis definition and regulatory pressure scoring

Four ceramide clearance axes were defined by manual curation of the KEGG sphingolipid metabolism pathway (hsa00600; (*28*)(*29*), selecting genes encoding enzymes that catalyze the committed step(s) of each clearance route: (1) the S1P axis (ACER1, ACER2, ASAH1, ASAH2, SPHK1, SPHK2); (2) the C1P axis (CERK); (3) the SM axis (SGMS1, SGMS2); and (4) the GSL axis (UGCG, UGT8, B4GALT4, B4GALT5). To estimate the net transcriptional drive toward each axis in each metacell, a regulatory pressure score was computed. For a given axis gene set G, the pressure on target gene g in metacell s is defined as P_gs = (Σ_r w_rg · A_rs) / (Σ_r |w_rg|), where the sum runs over all regulators r in the ARACNe3 network, w_rg = tfmode_rg × likelihood_rg is the signed regulatory weight, and A_rs is the VIPER activity of regulator r in metacell s. The axis-level score M_s is the mean of P_gs across all genes g in the axis gene set.

### Dominant axis classification

Each metacell was then classified according to its dominant axis, the axis receiving the highest regulatory pressure, provided that the margin over the second-ranked axis exceeded a defined threshold. Metacells with insufficient margin were classified as ambiguous. Each metacell was assigned a dominant sphingolipid axis based on its four pressure scores, subject to two empirically determined quality filters: the overall pressure range across four axes (maximum minus minimum) had to exceed the 25th percentile threshold (range4 ≥ 1.79), and the margin between the top-ranked and second-ranked axis had to exceed its 25th percentile threshold (margin4 ≥ 0.19). Metacells failing either criterion were classified as “Ambiguous.”

### Metabolic diversity quantification

At the single-metacell level, the four axis pressure scores were transformed to a probability distribution via the softmax function (β = 1.0), and Shannon entropy was computed as the per-metacell metabolic diversity index. At the population level, the four-dimensional pressure space was discretized into quantile bins (K = 6 bins per axis) and the joint Shannon entropy across all occupied bins was computed for each condition.

### Identification of candidate axis regulators using msVIPER

For each condition–axis combination, metacells were ranked by axis pressure score, and the top and bottom 20% were selected as high-and low-activity groups, respectively. A gene expression signature was computed using row-wise t-tests on the full-transcriptome CPM matrix, and the resulting z-scores were used as input to msVIPER (*59*) (minimum regulon size = 25). Candidate regulators were evaluated by integrating (1) msVIPER NES and Benjamini-Hochberg adjusted p-value (padj < 0.05); (2) axis gene linkage, defined as the sum of absolute regulatory weights connecting each regulator to the axis gene set; and (3) motif accessibility support from chromVAR, assessed by mapping each transcription factor to its cognate motif(s) in the JASPAR 2024 database. The top 20 regulators per axis were selected by mean msVIPER significance across conditions and visualized as dot plots encoding NES (color), axis gene linkage (dot size), and motif accessibility significance (transparency).

## Acknowledgements

We gratefully acknowledge funding from Physical Sciences Oncology Network at the National Cancer Institute (grant U01CA261841) and R01 grant R01CA272601. The authors wish to acknowledge the Biological Mass Spectrometry Shared Resource at the Stony Brook Cancer Center for expert assistance with running MALDI-MSI. We also appreciate the critical reading and feedback from Dr. Chiara Luberto.

## Supplementary Information

**Table 1**

Annotation for untargeted MALDI phenotyping

**Table 2**

Gene list for lipid CRISPR library

**Table 3**

Organoid Media Recipe

