## Extended Figures for "Metabolic plasticity enables cancer adaptation to acidic tumor ecosystems"

### Metabolic plasticity of sphingolipids governs cancer cell fitness in acidic tumor ecosystems

#### Abstract:

Cancer cells must adapt to harsh tumor microenvironments, including acidic stress, to survive and thrive. Understanding how cancer cells achieve this adaptation can uncover new biomarkers and therapeutic strategies. In this study, we investigated the spatial metabolic phenotypic heterogeneity of breast cancer cells in acidic habitats using spatial multi-omics approaches on 3D spheroids. We found cancer cells dynamically regulate sphingolipid metabolism to finetune their cell state to cope with acidic selection pressure. Cancer cells evolve mechanisms to deal with initially accumulating toxic ceramides but later adapt to it by rerouting SL metabolic pathways to eliminate them. Using advanced MALDI image analysis, and SL inhibitors on patient derived organoids, we demonstrated that cancer cells can switch between metabolic routes when key pathways are blocked, showcasing remarkable cell state plasticity. These insights highlight the potential to target metabolic plasticity as a novel therapeutic strategy to disrupt cancer adaptation and evolution, offering new avenues for cancer treatment.

**Key words:** Tumor ecology and evolution, Lipid phenotype, Acidic tumor microenvironment, Metabolic reprogramming, Sphingolipid signaling, Spatial phenotyping, Lipidomic, MALDI mass spectrometry

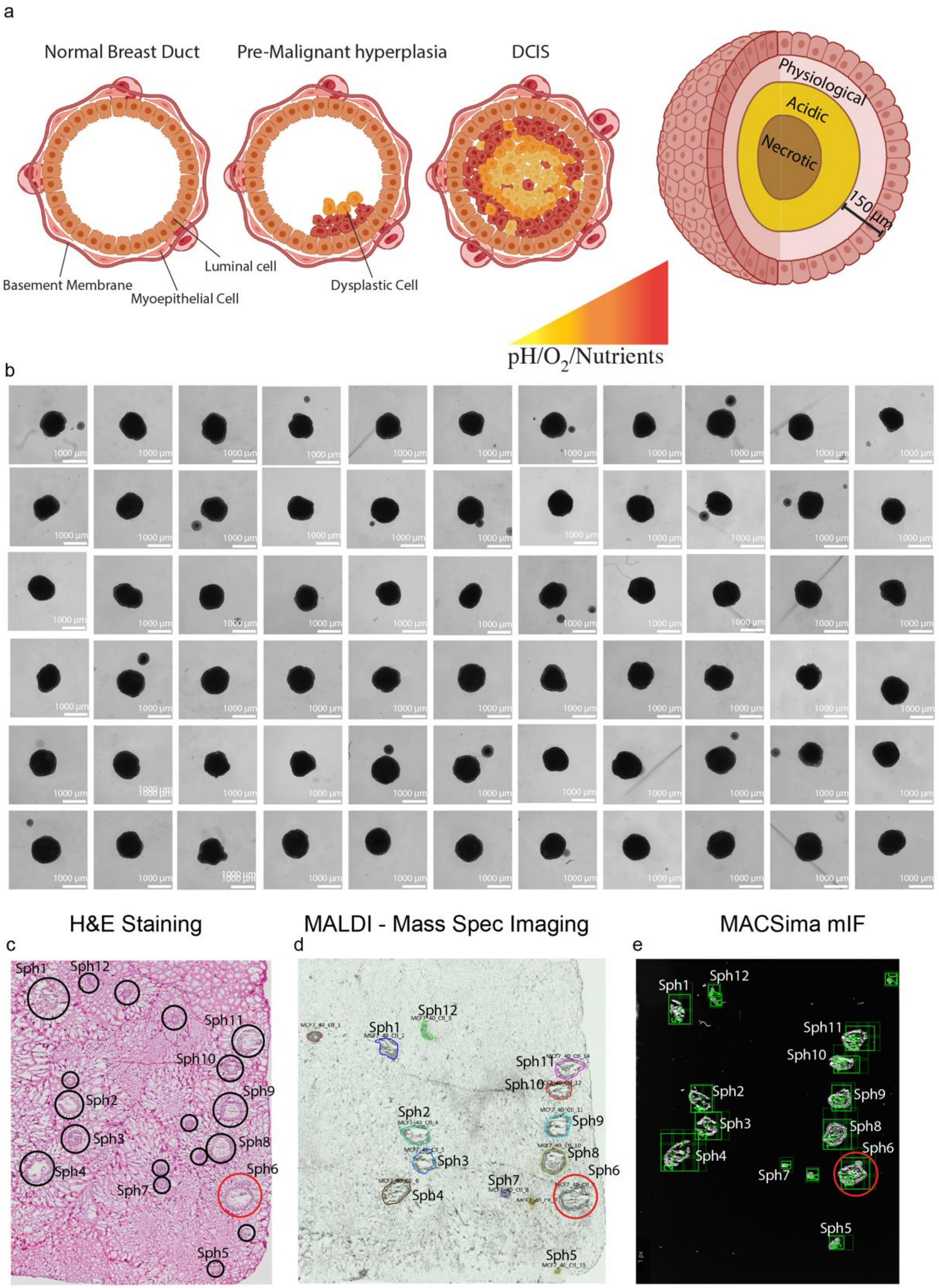

a
Normal Breast Duct
Pre-Malignant hyperplasia
DCIS
Physiological
Acidic
Necrotic
150 μm
Luminal cell
Basement Membrane
Myoepithelial Cell
Dysplastic Cell
pH/O2/Nutrients
b
1000 μm
H&E Staining
MALDI - Mass Spec Imaging
MACSima mIF
c
d
e
Sph1
Sph2
Sph3
Sph4
Sph5
Sph6
Sph7
Sph8
Sph9
Sph10
Sph11
Sph12

**Extended Figure 1. Spatial multi-omics framework for profiling acidic tumor ecosystems. a,** Schematic of breast tumor acidic TME in DCIS (left) and spheroids (right), highlighting gradients in pH, oxygen, and nutrient availability. **b,** 3D breast cancer spheroid images used for one spatial experiment and analyses. **c,** H&E-staining of the embedded spheroids in figure (a) and sections showing preserved spheres architecture. Red circles denote the spheroids shown in Figure 1. **d,** Sequential spheroid sections mounted on conductive slides with annotated regions of spheres for MALDI imaging. Red circles denote the spheroids shown in Figure 1. **e,** Representative overview image acquired by the MACSima multiplex immunofluorescence platform, used for spatial registration and region selection of the sequential cut of the same block in b and c. Red circles denote the spheroids shown in Figure 1. Spatial registration of MALDI lipid ion maps with corresponding H&E and multiplexed imaging, enabling single-spheroid–resolved metabolic and cellular profiling.

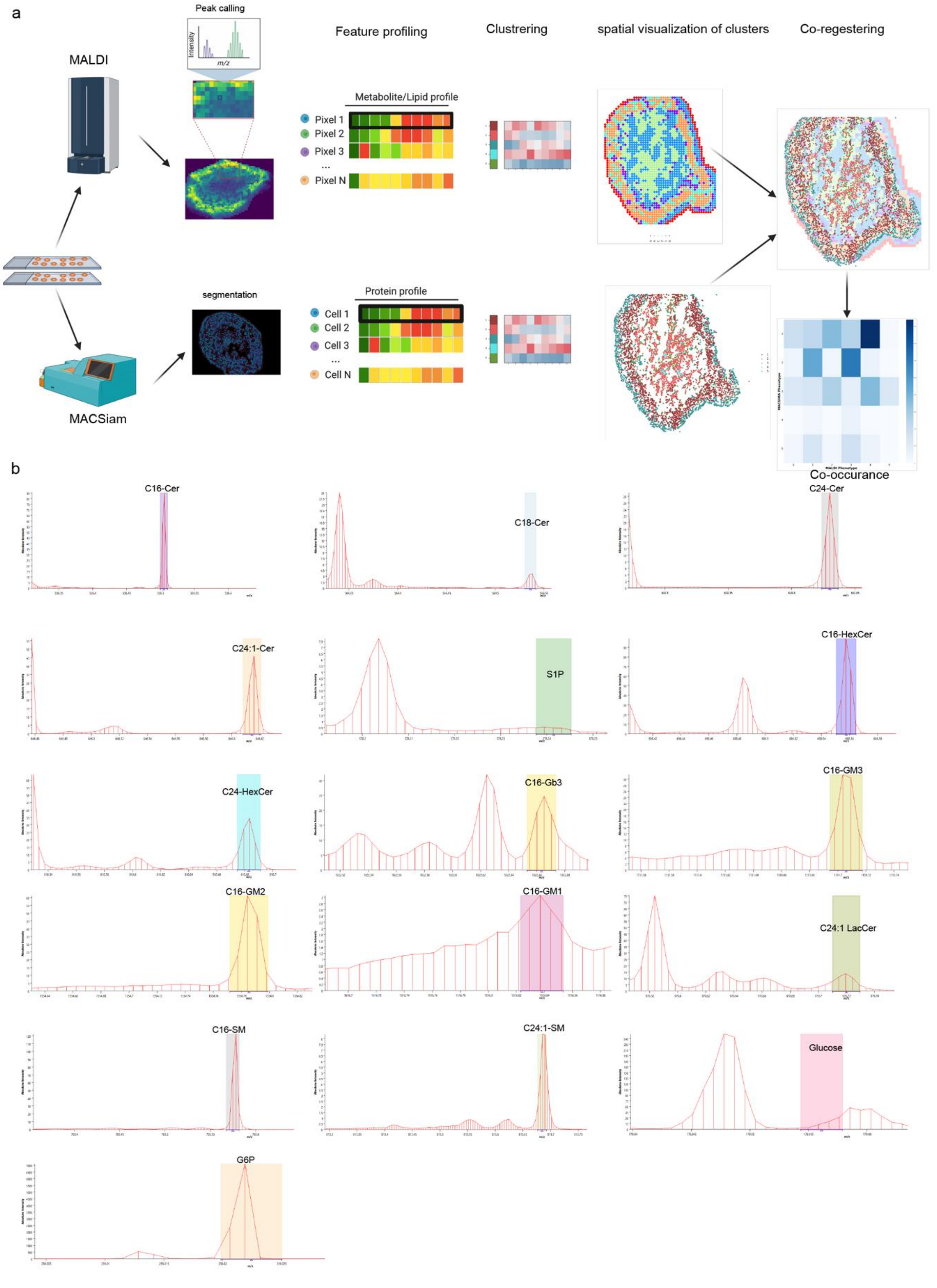


**Extended Figure 2. MALDI-MSI and MacSIMA-mIF spatial analysis reveals ecological patterns in MCF7 Spheroids.** **a**, schematic of the spatial data analysis pipeline: Sequential sample sections were analyzed using Matrix-Assisted Laser Desorption/Ionization (MALDI) mass spectrometry for metabolomics and MACSima imaging for proteomics. MALDI imaging generated a spatial grid of 20 × 20 µm pixels, with each pixel containing spectral peaks corresponding to hundreds of molecular species. Metabolite annotation was performed using a customized feature list or MetaboSpace. A two-step clustering algorithm was employed to categorize pixels with similar metabolomic profiles, which were subsequently visualized on the original sample. For MACSima-based proteomics, single-cell segmentation was performed to extract individual cellular protein expression profiles, generating a matrix where each cell was characterized by its expression levels for each protein marker. Analogous to the MALDI workflow, clustering was applied to group cells based on their proteomic signatures and visualized within the sample. To integrate these datasets, the metabolomic clusters from MALDI and the proteomic clusters from MACSima were manually co-registered, allowing each cell to retain its proteomic phenotype while also being assigned a corresponding metabolomic signature. The spatially co-registered dataset was then used to calculate the co-occurrence of metabolic phenotypes with proteomic phenotypes. **b**, Representative MALDI peak spectra for selected metabolites and lipids, illustrating the characteristic m/z peaks associated with key molecules detected in the samples. The peak intensities represent the average spectral signal for each identified metabolite.

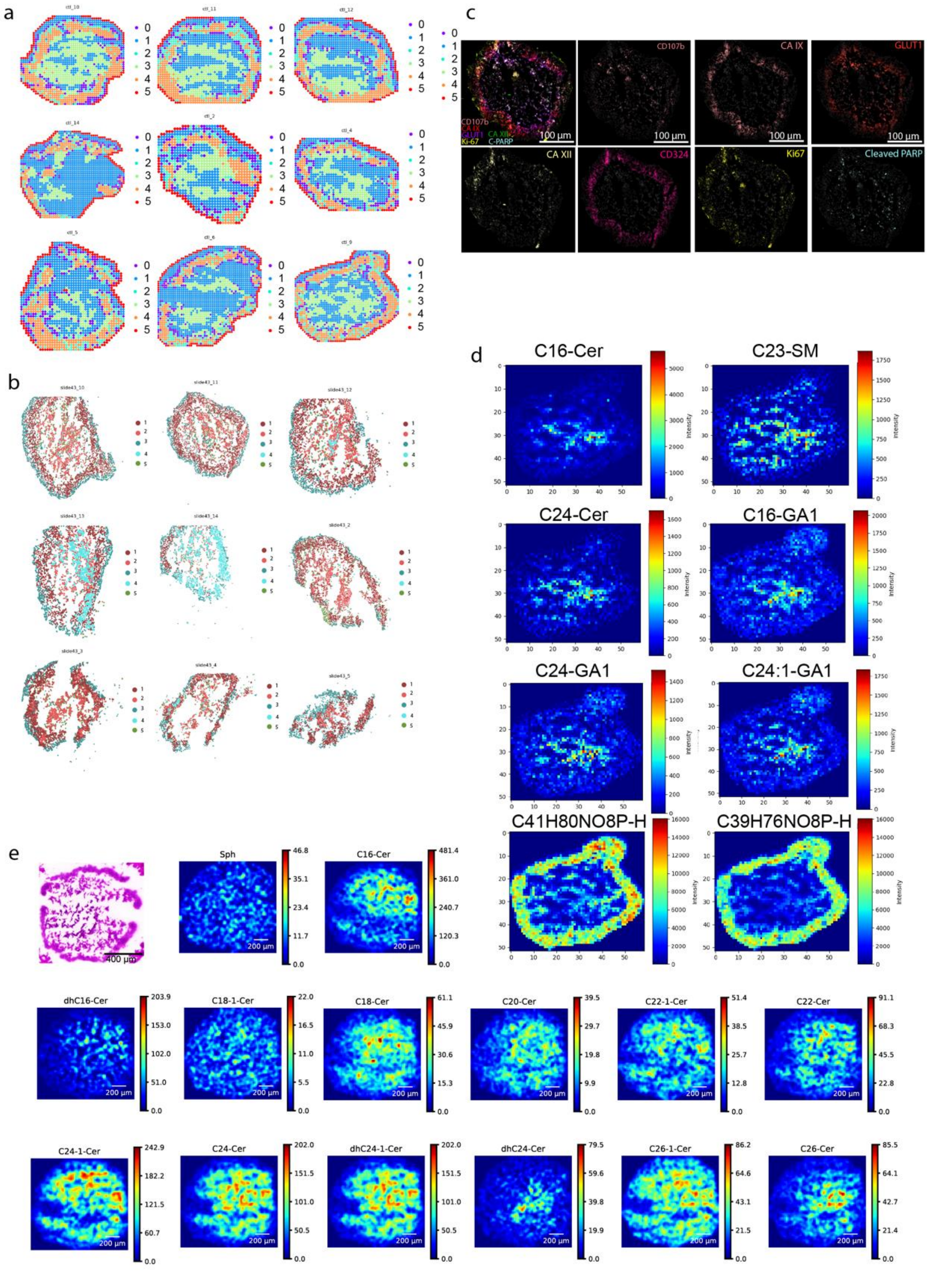

**Extended Figure 3. Registration of MALDI-MSI and MACSima-mIF spatial data reveals sphingolipid co-localization in acidic niches.** a, Unsupervised metabolic clustering of multiple spheroids. b, Unsupervised clustering of proteomic data, illustrating the spatial organization of proteomic phenotypes across multiple spheroids. c, Multiplex immunofluorescence (mIF) imaging using MACSima, visualizing key microenvironmental markers (CD107b, CA IX, Glut1, Ki-67, CA XII, cPARP, CD324) to define phenotypic niches within different tumor microenvironments. d, Spatial intensity maps depicting the distribution of sphingolipids enriched in cluster-3 and lipids enriched in cluster-4. e, Spatial intensity maps of selected Ceramides.

#### Extended Figure 4

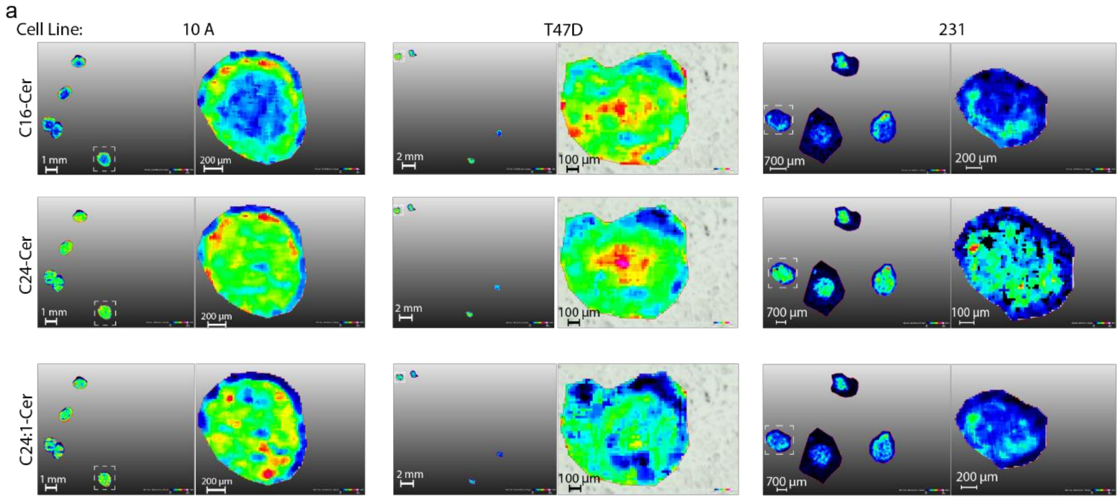


**Extended Figure 4. Ceramides are upregulated in the acidic niches of breast cancer spheroids.** MALDI-MSI spatial heatmaps in a, MCF10A , b, T47D , and c, MDA-MB231 spheroids of various ceramide species.

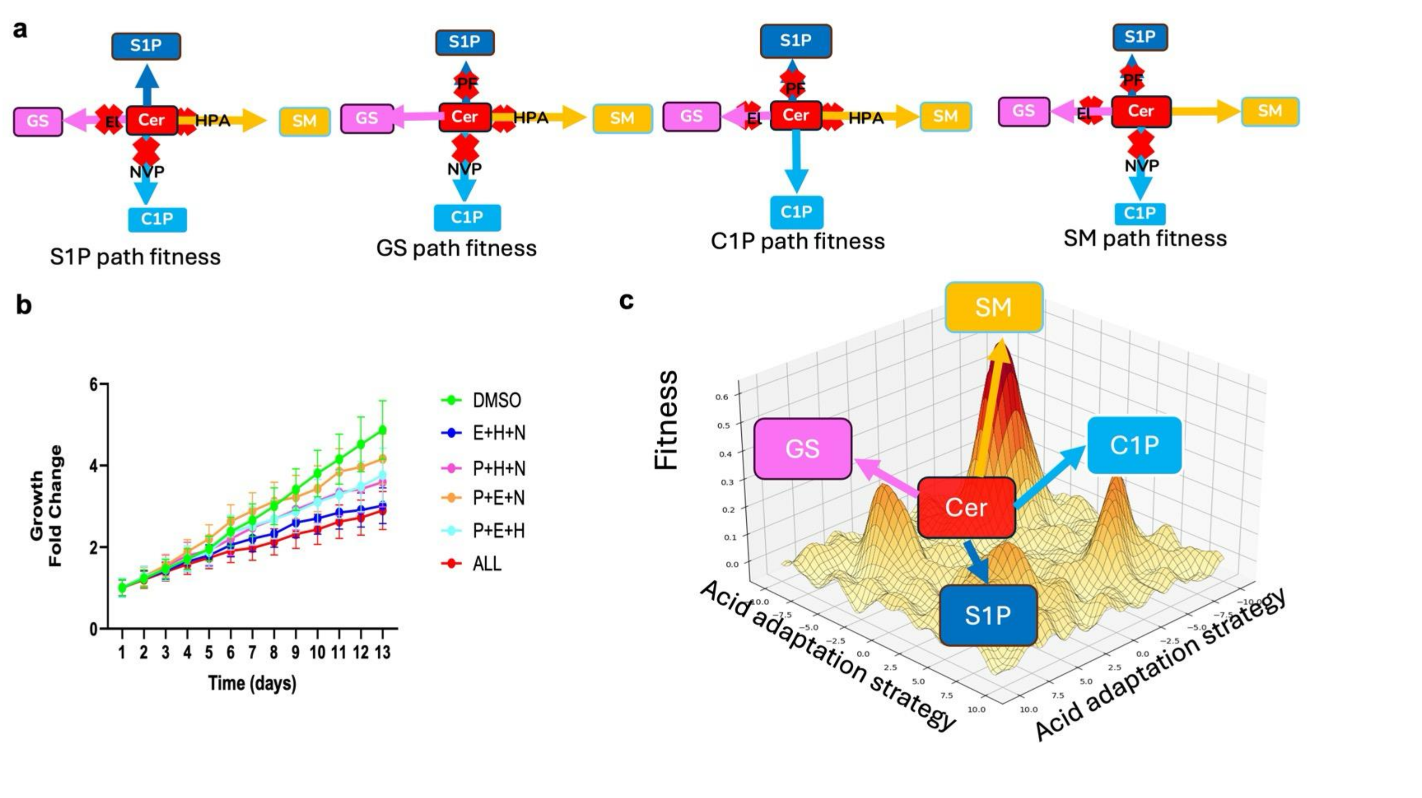


**Extended Figure 5. Fitness landscape of ceramide clearance pathways under chronic acidosis.**
**a,** Schematic representation of ceramide clearance strategies in which three of the four major sphingolipid pathways are pharmacologically inhibited, leaving a single pathway (S1P, GS, C1P, or SM, respectively) available for ceramide disposal. Red crosses denote inhibited enzymatic nodes, enabling direct assessment of the fitness contribution of each individual ceramide clearance route. **b,** Growth kinetics of MCF7 spheroids under leave-one-pathway–available conditions, demonstrating differential fitness depending on the remaining ceramide clearance route. **c,** Conceptual fitness landscape illustrating ceramide as a central metabolic hub and individual sphingolipid fates as distinct adaptive strategies. Peaks in fitness correspond to pathways that more effectively support growth under chronic acidosis, highlighting metabolic degeneracy and plasticity in ceramide handling.

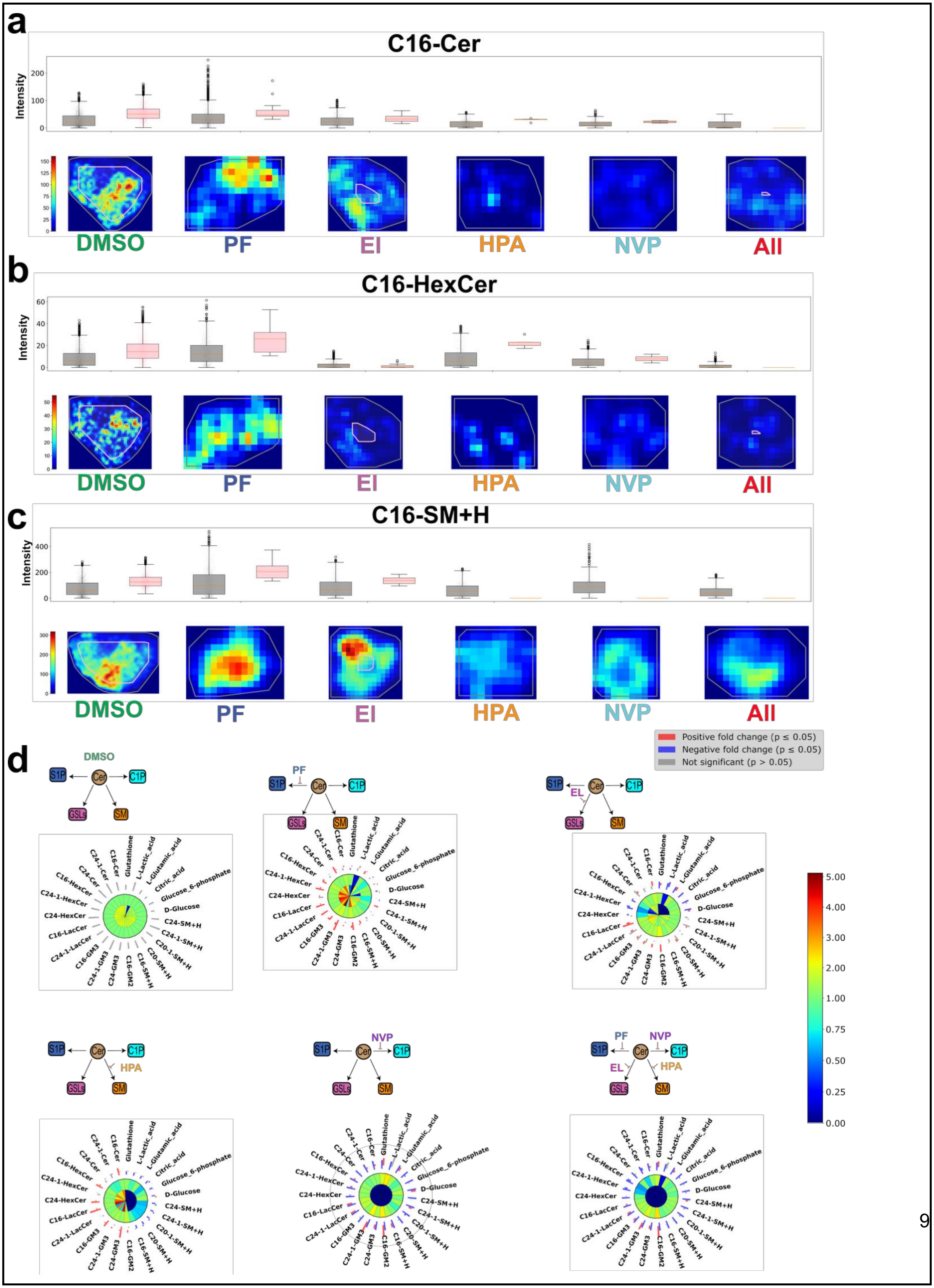
a
C16-Cer
Intensity
b
C16-HexCer
c
C16-SM+H
DMSO
PF
EI
HPA
NVP
All
d
Positive fold change (p ≤ 0.05)
Negative fold change (p ≤ 0.05)
Not significant (p > 0.05)
S1P
Cer
C1P
GSLs
SM
EL
C24-1-Cer
C16-Cer
C24-Cer
Glutathione
L-Lactic_acid
L-Glutamic_acid
Citric_acid
Glucose_6-phosphate
D-Glucose
C24-SM+H
C24-1-SM+H
C20-1-SM+H
C20-SM+H
C16-SM+H
C16-GM2
C24-GM3
C24-1-GM3
C16-GM3
C24-1-LacCer
C16-LacCer
C24-HexCer
C24-1-HexCer
C16-HexCer

**Extended Figure 6. SPHERE analysis reveals pathway-specific sphingolipid remodeling in patient-derived organoids (PDOs). a–c,** Boxplots (top) and corresponding spatial ion maps (bottom) showing the abundance and spatial distribution of representative C16 sphingolipid species in PDOs, including C16-ceramide (C16-Cer), C16-hexosylceramide (C16-HexCer), and C16-sphingomyelin + $H^+$ (C16-SM+H), as measured by spatial lipidomics. **d,** SPHERE plots integrating fold-change directionality and statistical significance across sphingolipid species and associated metabolic intermediates under pathway-specific perturbations. Each sector represents a metabolite, with color indicating positive (red), negative (blue), or non-significant (gray) changes ($p \leq 0.05$). Schematic overlays illustrating the inferred redistribution of ceramide flux into distinct sphingolipid pathways, including S1P, glycosphingolipids (GSLs), sphingomyelin (SM), and ceramide-1-phosphate (C1P), under selective pathway inhibition in PDOs. Collectively, these analyses demonstrate heterogeneous yet coordinated sphingolipid plasticity in patient-derived tumor organoids, supporting ceramide clearance as a conserved adaptive strategy in both complex spheroid and organoid 3D tumor ecosystems.

**a**

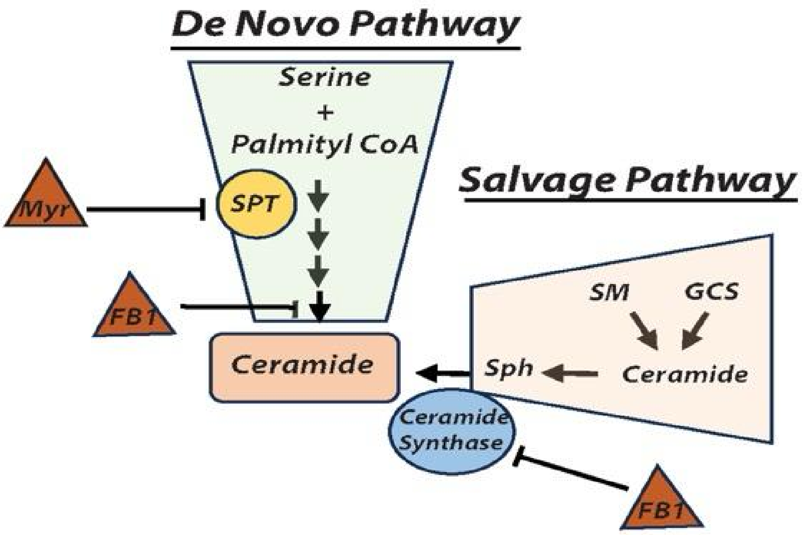


**b**

| Condition | Reduction by Myrocin(%) | Reduction by FB1 (%) | Difference (FB1 - Myrocin) (%) |
|---|---|---|---|
| NA 6.5 | 38.57 | 64.45 | 25.88 |
| AA 6.5 | 61.85 | 82.35 | 20.50 |

De novo pathway contribution

Salvage pathway contribution

**Extended Figure 7. Differential contribution of de novo and salvage pathways to ceramide production under acidic conditions. a,** Schematic representation of ceramide biosynthesis through the de novo pathway, initiated by serine palmitoyltransferase (SPT), and the salvage pathway, in which ceramide is regenerated from sphingomyelin (SM) and glycosphingolipids (GCS). Pharmacologic inhibitors myriocin (Myr) and fumonisin B1 (FB1) are shown at their respective enzymatic targets. **b,** Quantitative analysis of ceramide reduction following inhibition of the de novo (myriocin) or salvage (FB1) pathways in non-acid-adapted NA- and AA-MCF7 cells cultured at pH 6.5. The relative difference between FB1- and myriocin-mediated reductions highlights the increased contribution of the salvage pathway to ceramide production under chronic acidosis.

**Extended Figure 8:**

MALDI mass spectrometry imaging (MALDI-MSI) of MCF7 spheroids treated with SL inhibitors, highlighting spatial distribution and intensity of selected molecules. Box plots indicate average molecular intensities in inner and outer layers, defined by a 125 μm threshold from the spheroid boundary.

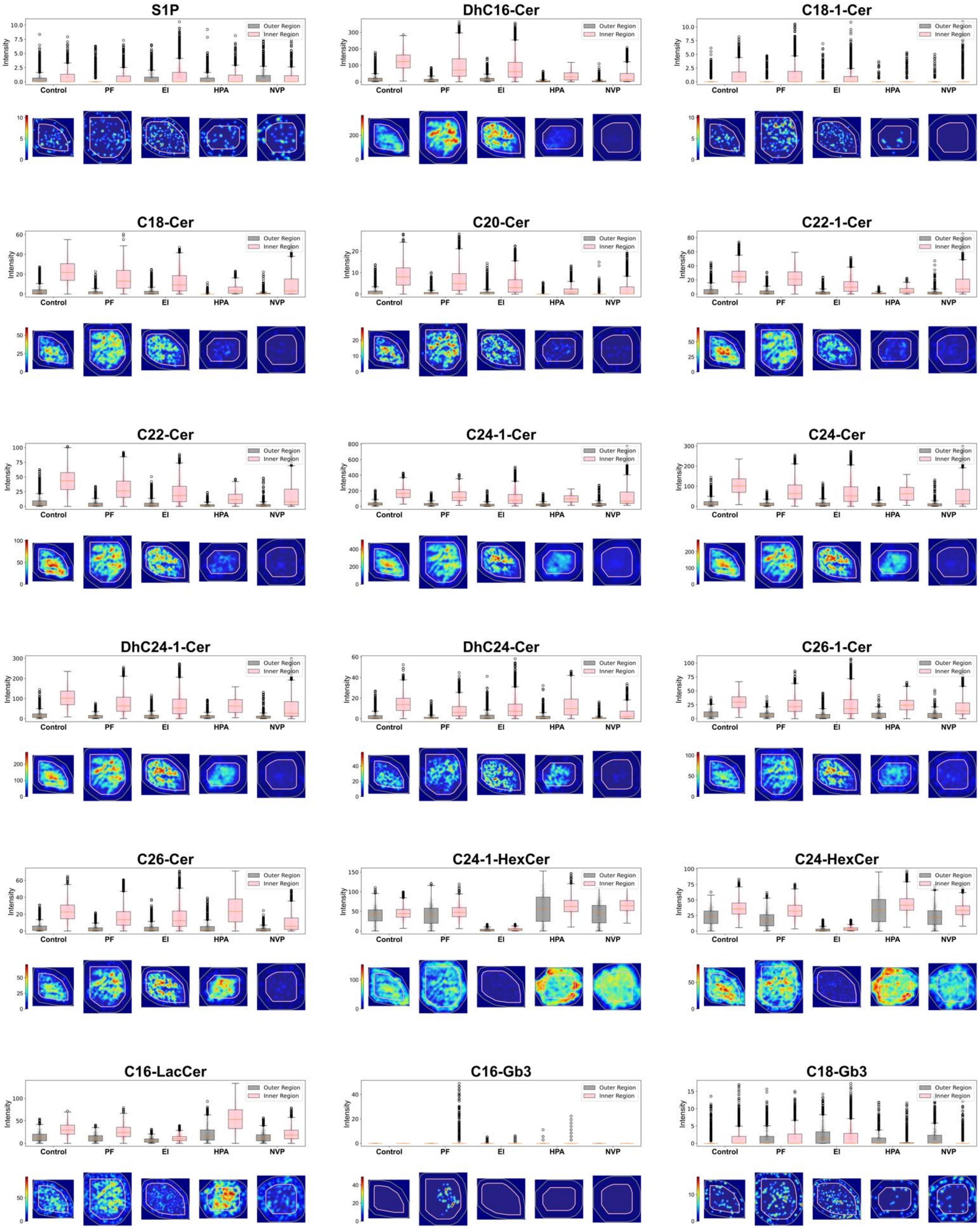

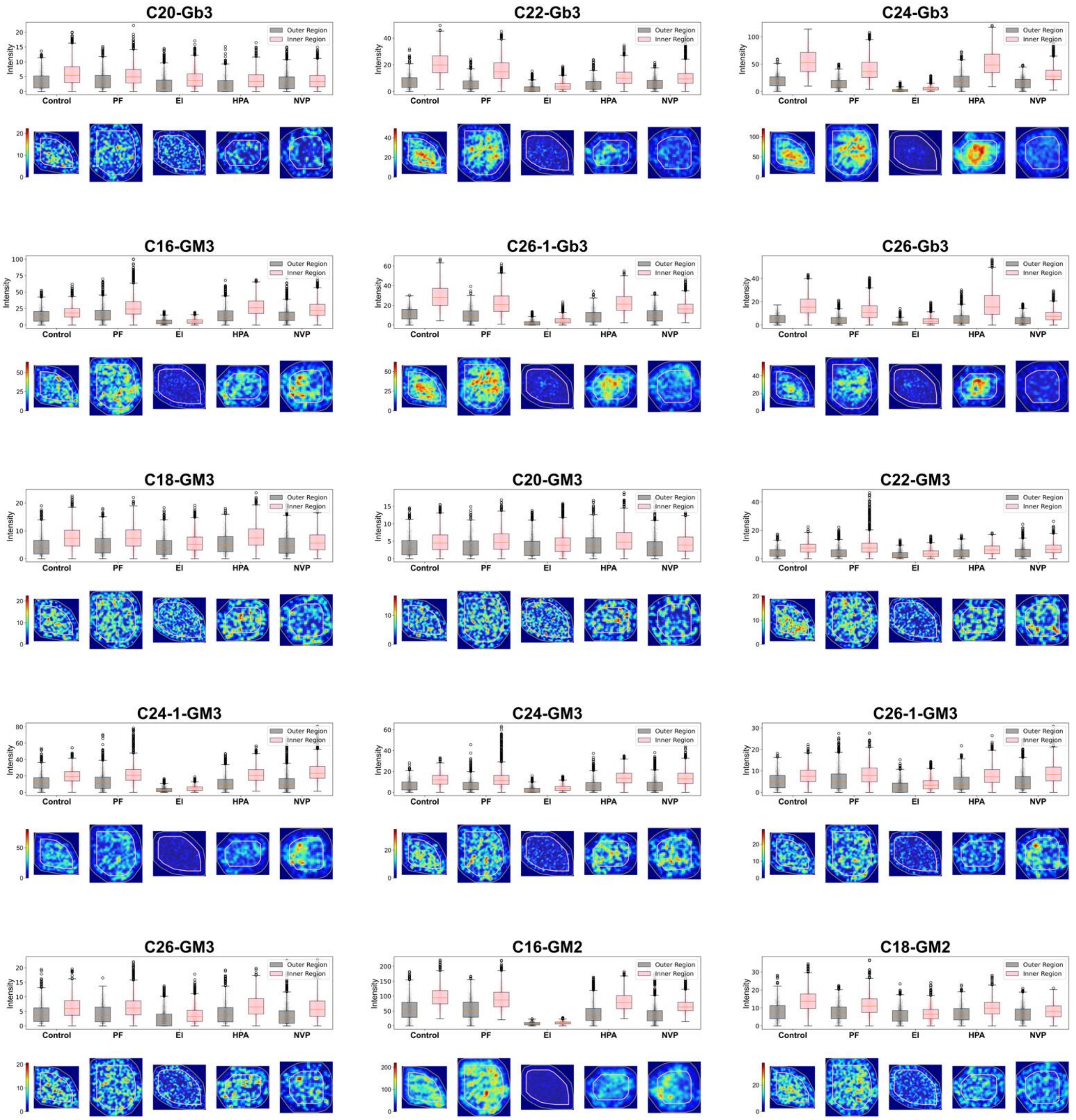
C20-Gb3
C22-Gb3
C24-Gb3
C16-GM3
C26-1-Gb3
C26-Gb3
C18-GM3
C20-GM3
C22-GM3
C24-1-GM3
C24-GM3
C26-1-GM3
C26-GM3
C16-GM2
C18-GM2
Outer Region
Inner Region
Intensity
Control
PF
EI
HPA
NVP

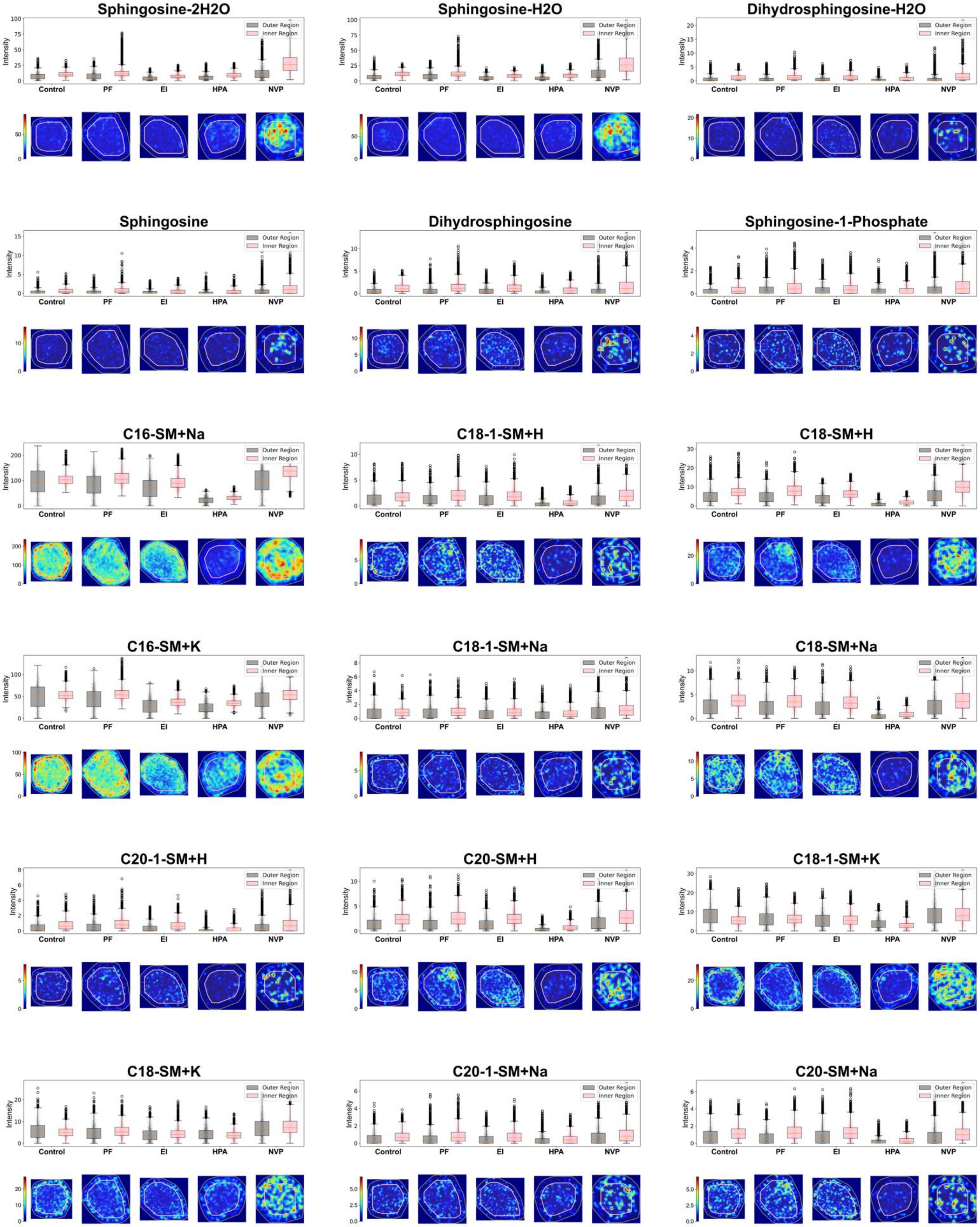


### Supplementary Information:

**Table1**

**Annotation for untargeted MALDI phenotyping**

#### Table2

Gene list for lipid CRISPR library
